# Metformin preconditioning protects against myocardial stunning and preserves mitochondrial dynamics and protein translation in a mouse model of cardiac arrest

**DOI:** 10.1101/2021.08.24.457506

**Authors:** Cody A. Rutledge, Claudia Lagranha, Takuto Chiba, Kevin Redding, Donna B. Stolz, Eric Goetzman, Sunder Sims-Lucas, Brett A. Kaufman

**Author notes:** **Corresponding Author:** Brett A. Kaufman, 200 Lothrop Street, BST E1241, Pittsburgh, PA 15261, 1-412-624-8644.

## Abstract

Cardiac arrest (CA) causes high mortality due to multi-system organ damage attributable to ischemia-reperfusion injury. Recent work in our group found that among diabetic patients who experienced cardiac arrest, those taking metformin had less evidence of cardiac and renal damage after cardiac arrest when compared to those not taking metformin. To investigate the potential for metformin to impact cardiac arrest outcomes, the current study investigates metformin interventions on cardiac and renal outcomes in a non-diabetic CA mouse model. We found that two weeks of metformin pretreatment protects against reduced ejection fraction and reduces kidney ischemia-reperfusion injury at 24 hours post-arrest. This cardiac and renal protection depends on AMPK signaling, as demonstrated by outcomes in mice pretreated with the AMPK activator AICAR or metformin plus the AMPK inhibitor compound C.

At this 24-hour time point, gene expression analysis showed that metformin pretreatment caused changes supporting autophagy, antioxidant response, and protein translation. Further investigation found associated improvements in mitochondrial structure and markers of autophagy. We also found that markers of protein nitrosylation between sham and arrest were not significantly different and thus unlikely to be a driver of gene expression differences. Notably, Western analysis indicated that protein synthesis was preserved in arrest hearts of animals pretreated with metformin. The AMPK activation-mediated protection of protein synthesis was also demonstrated in a hypoxia/reoxygenation cell culture model. Despite the positive impacts of pretreatment *in vivo* and *in vitro*, metformin did not preserve ejection fraction when deployed at resuscitation. Taken together, we propose that metformin’s *in vivo* cardiac preservation occurs through AMPK activation, requires adaptation before arrest, and is associated with preserved protein translation.

**New and Noteworthy:** We leveraged a mouse model of cardiac arrest to study whether metformin pretreatment could protect cardiac function after arrest. We find that AMPK activity is crucial to this protection and characterize several pathways associated with this response. Signaling nodes with strong association may be precise targets to promote cardiac function in ischemia/reperfusion injury or in cardiac stunning.

## Introduction

Cardiac arrest (CA) refers to the abrupt cessation of cardiac function and affects over 600,000 patients annually in the United States (1,2). Patients who return spontaneous circulation (ROSC) after CA experience systemic ischemia-reperfusion injury, typically resulting in multi-system organ damage. Typical findings include cardiogenic shock, acute renal failure, liver damage, and neurologic dysfunction (3–5). Previous observational studies in cardiac arrest patients have shown that low cardiac ejection fraction (EF) (6) and reduced kidney function (7,8) after arrest are predictors of increased mortality. Despite the prevalence of CA, no pharmacologic therapy has been shown to improve overall survival.

A recent study by our group found that diabetic patients taking metformin had less evidence of cardiac and renal damage after CA than diabetic patients on other treatments (9), suggesting that metformin pretreatment may be beneficial to CA patients through an undefined mechanism. Metformin is an oral antihyperglycemic agent used as a first-line agent for type 2 diabetes that enhances insulin sensitivity and normalizes glucose and lipid homeostasis (10–13). Beyond its role in controlling diabetes, metformin has demonstrated clinical benefit across a wide variety of pathologies, including decreased mortality in the setting of coronary artery disease (14), congestive heart failure (15), acute kidney injury (16), chronic kidney disease (15), septic shock (17,18), and major surgical procedures (19). Cardiovascular studies suggest that improved outcomes occur independently of the glucose-lowering effects of metformin and may instead be attributable to metformin’s pleiotropic effects (20). In a preclinical rat model of CA, metformin pretreatment improved neurologic outcomes and survival (21), and similar protection was recently reported using metformin as a post-arrest therapy (22). However, the mechanisms underlying this protection and the value of metformin as a rescue therapy after CA remain unclear.

Several mechanisms beneficial to cardiovascular health have been implicated in metformin’s effects, including reduced oxidative stress, anti-apoptotic activities, JNK inhibition, complex I inhibition, and AMP Kinase (AMPK) activation (21,23,24). AMPK activation is prevalent in the metformin literature, and AMPK activity is essential for survival after ischemic stress (25), but it is unclear whether AMPK activity is the mediator of metformin-mediated protection in CA. Furthermore, it is unclear whether the adaptive upregulation of AMPK during cardiac stress is optimal or whether further activation could even more strongly impact recovery in cardiac injury models.

In this study, we sought to test metformin’s potential benefit on heart and kidney protection after CA and begin to assess the drivers of the effects witnessed clinically in our earlier diabetic patient study. We evaluated cardiac and renal outcomes one day after CA in untreated sham (Sham), untreated arrest (Arrest), metformin-pretreated sham (Sham+Met), and metformin pretreated arrest mice (Arrest+Met) mice. Additionally, we tested the AMPK-dependence of metformin’s effects by pretreating with the AMPK activator 5-aminoimidazole-4-carboxamide-1-β-D-ribofuranoside (AICAR; Arrest+AICAR) instead of metformin or by inhibition of AMPK via compound C given in the presence of metformin (Arrest+Met+Comp C). We also evaluated a small cohort of mice with intravenous metformin as rescue therapy. From the pretreatment group, we identified transcriptomic changes in metformin-treated sham and arrest mice and tested many of the pathways implicated by Western blot analysis or confirmed changes in a cell model of hypoxia/reoxygenation (H/R).

## Methods

### Sudden Cardiac Arrest Model

Eight-week-old male and female C57BL/6J mice (Jackson Labs, Bar Harbor, ME, #000664) underwent cardiac arrest by delivery of potassium chloride (KCl) directly into the LV by percutaneous, ultrasound-guided needle injection (Figure 1) as previously described (26). Briefly, mice were anesthetized using vaporized isoflurane (Henry Schein, Melville, NY, #1182097) and endotracheally intubated and mechanically ventilated (MiniVent, Harvard Apparatus, Holliston, MA, #73-0043) at 150 bpm and volume of 125 μL for females and 140 μL for males. Body temperature was maintained using a rectal temperature probe and heating pad (Indus Instruments, Webster, TX, #THM150). The chest was cleaned of hair using Nair and sterilized with betadine before the introduction of a 30-gauge needle into the LV under ultrasound guidance (Visual Sonics Vevo 3100 with Vevo LAB v 5.5.1 software, Toronto, Canada), followed by delivery of 40 μL of 0.5M KCl to induce asystole. The ventilator was stopped, and the mice remained in asystole for 8 minutes. 7.5 minutes after KCl dosing, 500 μL of 15 μg/mL epinephrine in saline (37 °C) was injected into the LV over approximately 30 seconds. At 8 minutes, the ventilator was restarted, and CPR was initiated by finger compression at about 300 bpm for 1-minute intervals. An electrocardiogram (ECG) was evaluated for return of sinus rhythm after each 1-minute interval. Animals not achieving ROSC by 3 minutes after CPR initiation were euthanized. Mice remained on ventilator support until spontaneous breathing frequency was greater than 60 times per minute. Sham mice received no KCl, but rather a single injection of epinephrine. After the procedure, all animals were placed in a recovery cage under a heat lamp.

**Figure 1.**
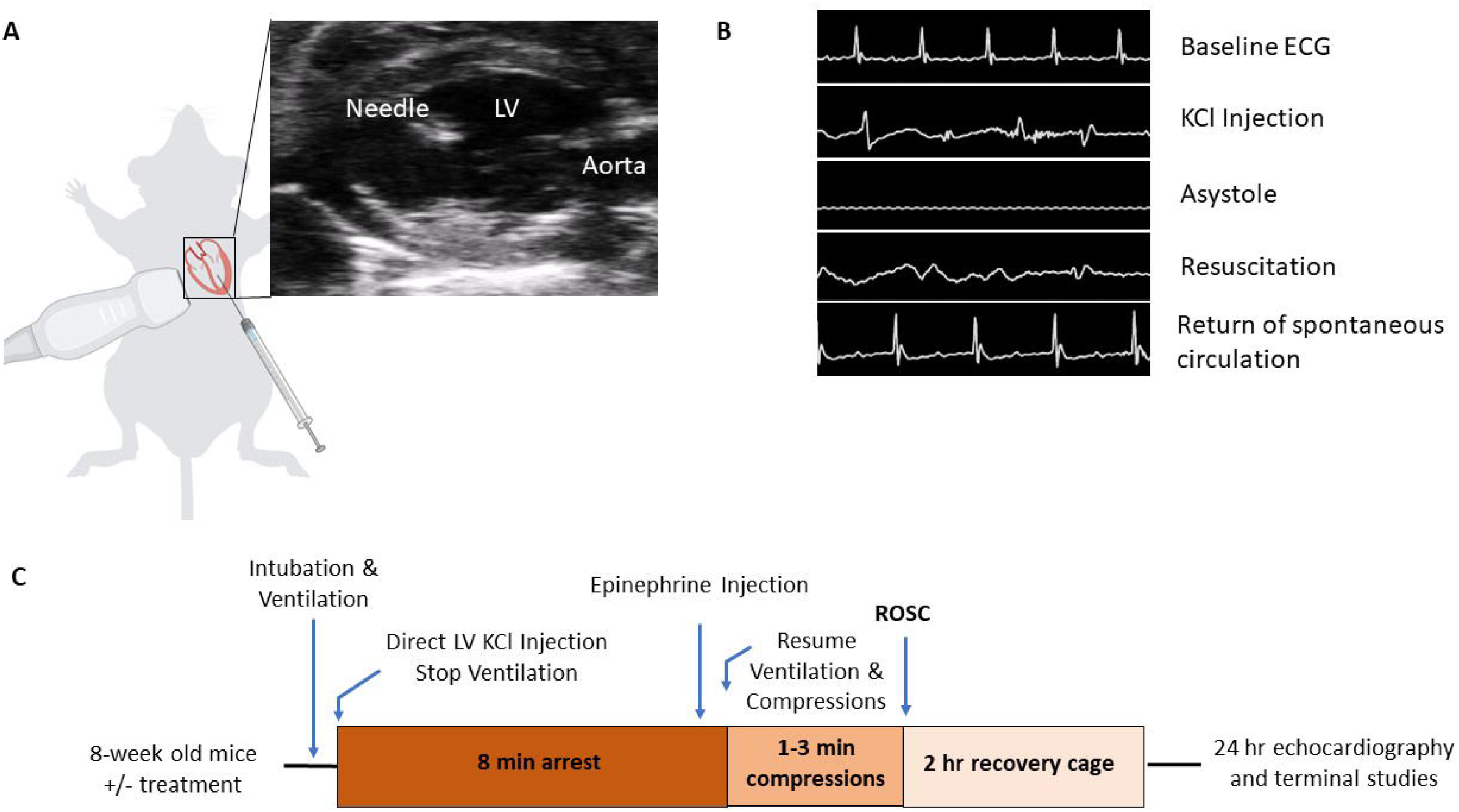
Mouse model of cardiac arrest (CA). A) Cartoon representation of direct left ventricular injection of potassium chloride (KCl) to cause asystole with representative ultrasound image of needle guidance. B) Representative electrocardioagrm (ECG) traces throughout time course of cardiac arrest procedure C) Time course of CA protocol. LV, left ventricle; ROSC, return of spontaneous circulation.

### Animal Treatment Groups

Treatment groups in the study were as follows: untreated sham (Sham), untreated cardiac arrest (Arrest), metformin-pretreated sham (Sham+Met), metformin-pretreated cardiac arrest (Arrest+Met), 5-aminoimidazole-4-carboxamide-1-beta-D-ribofuranoside (AICAR) pretreated cardiac arrest (Arrest+AICAR), metformin + compound c pretreated cardiac arrest (Arrest+Met+Comp C), and metformin rescue treatment cardiac arrest (Arrest+Rescue Metformin). Metformin pretreatment consisted of *ad libitum* access to metformin (Major Pharmaceuticals, Livonia, MI, #48152) in drinking water (1 mg/mL) for 14 days before the procedure. AICAR pretreated mice were given IP injections of 500 mg/kg AICAR (Toronto Research Chemicals, Toronto, CA, #A611700) in saline every other day for 14 days before CA. Compound C pretreated mice were given 20 mg/kg IP injections of compound C (Cayman Chemical, Ann Arbor, MI, #11967) in saline daily for 14 days before arrest. Metformin rescue therapy was given as a single direct LV injection (1250 μg/kg) dissolved in saline and 500 μL of 1 mg/mL epinephrine (Par Pharmaceutical, Chestnut Ridge, NY, #10977) at the time of resuscitation. Mice were randomly assigned to treatment groups. Male and female mice were used in all groups. All mouse studies were performed at the University of Pittsburgh in compliance with the National Institutes of Health Guidance for Care and Use of Experimental Animals. This protocol was approved by the University of Pittsburgh Animal Care and Use Committee (Protocol #18032212).

### Echocardiography and Ultrasound

Immediately before arrest CA, mice were evaluated by transthoracic echocardiography using Vevo 3100 imaging systems (Visual Sonics) with a 40 MHz probe as previously described (27). Repeat echocardiography was performed one day after arrest under isoflurane anesthesia delivered by nose cone. Heart rate was maintained between 400-500 bpm during imaging by adjusting isoflurane concentration. B-mode images taken from the parasternal long-axis were captured, and LV EF was calculated using modified Simpson’s methods (28). A cohort of all groups was assessed for renal perfusion after resuscitation. The ultrasound probe was oriented transversely across the abdomen at the plane of the right kidney and monitored for renal artery blood flow by Doppler imaging. All image analysis was performed by a blinded sonographer (Vevo Lab 5.5.1, Visual Sonics).

### Tissue and Serum Collection

After euthanasia with isoflurane and cervical dislocation, mice underwent cardiac puncture to collect blood by heparinized syringe. Blood was separated by centrifugation at 2,000 x g at 4°C for 10 minutes, and the serum was flash frozen. These samples were evaluated for blood urea nitrogen (BUN) and creatinine by the Kansas State Veterinary Diagnostic Laboratories (Manhattan, KS). Hearts from the mice were excised, and LVs were isolated and flash frozen.

### Western Blot

Frozen LV tissue was homogenized in lysis buffer containing a protease/phosphatase cocktail (Sigma-Aldrich, St. Louis, MO, #11697498001) and normalized for protein content using a BCA assay (Life Technologies, Carlsbad, CA, #23235). Samples were separated on NuPage 4-12% gradient SDS-PAGE gels (ThermoFisher, Waltham, MA, #WG1403BOX) and iBlot transferred onto nitrocellulose membranes (Invitrogen, #IB21001). Membranes were blocked in 5% milk (non-phosphorylated antibodies) or 5% BSA (phosphorylated antibodies) for 1 hour and then incubated overnight at 4 °C with primary antibodies, including, p-AMPK (Thr172, 1:1000, Cell Signaling, Danvers, MA, #2535), AMPK (1:1000, Cell Signaling, #2532), GAPDH (1:5000, Millipore, St. Louis, MO, #AB2302), p62 (1:1000, Sigma-Aldrich, #P0067), LC3 (Invitrogen, 1:1000, PA1-16931), OxPhos Rodent Antibody Cocktail 1:5000 (ThermoFisher, #458099), OPA1 (Invitrogen, 1:1000, PA5-57875), MFN2 (Invitrogen, 1:1000, #711803), p-DRP1 (Ser616, 1:1000, Thermo, #PA5-64821), DRP1 (1:1000, Cell Signaling, #5391), p-S6 (Ser240/244, 1:1000, Cell Signaling, #2215), S6 (1:1000, Cell Signaling, #2217), p-4EBP1 (Thr37/46, 1:1000, Cell Signaling, #2855), 4EBP1 (1:1000, Cell Signaling, #9644), nitrotyrosine (Santa Cruz, 1:1000, #32757) and ATF4 (1:1000, Thermo, #PA5-27576). Following incubation, membranes were washed with TBS-tween and then probed for 1 hour at room temperature with anti-mouse or anti-rabbit secondary antibodies conjugated with either horse radish peroxidase (Jackson ImmunoResearch, West Grove, PA, #115-035-003 and #115-035-144) or infrared secondary (IRDye 680 and 800, LICOR, Lincoln, NE, #926-32210 and #926-68071). Images were obtained by developing on a ChemiDoc XRS imaging system (BioRad, Hercules, CA) or Odyssey CLx infrared imaging system (LICOR) and analyzed using ImageJ software (National Institutes of Health, Bethesda, MD).

### Microarray Analysis

Microarray analysis was performed on cDNA through the Affymetrix microarray analysis service (ThermoFisher). Sham, Arrest, Sham+Met, and Arrest+Met mice were included in these studies with an even distribution of males and females (total n=6 per group). Differential gene expression analysis was performed using Transcription Analysis Console (Thermofisher). Gene-level p-values less than 0.05 were considered significant for gene inclusion. Subsequent pathway analysis was performed to compare untreated sham and arrest groups through Ingenuity Pathway Analysis (Qiagen). Complete datasets were deposited in GEO (accession no. GSE176494).

### Cell Model

AC16 cells, a hybrid cell model derived from the fusion of adult ventricular myocytes with stable, proliferating SV40 fibroblasts (29), were cultured in DMEM with 10% FBS and penicillin/streptomycin (Thermo, #15140122). Cells were incubated overnight at 37° C in DMEM with and without AICAR (1.25 mM) before the hypoxia/reoxygenation (H/R) challenge. Immediately prior to challenge, media was replaced with either DMEM (normoxia group) or Esumi buffer containing 137 mM NaCl, 12 mM KCl, 0.5 mM MgCl_2_, 0.9 mM CaCl_2_, 20 mM HEPES, and 20 mM 2-deoxy-d-glucose at pH 6.2 (H/R group). Cells were challenged with either normoxia (20% oxygen, 5% CO_2_) or hypoxia (1% oxygen, 5% CO_2_) for 4 hours, and then media was exchanged with normoxic DMEM. After 30 min of normoxia, cells were stained with NucBlue Live (Hoechst 33342, Invitrogen) and NucGreen Dead (Sytox green, Invitrogen) according to manufacturer recommendations and visualized on a Cell Insight CX7 HCX Platform (Thermo) at 20x to quantify the ratio of living to dead cells. These cells were pelleted and flash frozen for protein quantification.

### Protein Synthesis Assay

The normoxia vs. H/R challenge was repeated for protein synthesis assays using the methionine analog 1-azidohomoalanine (AHA; Thermo). Immediately after normoxia/hypoxia, cell media was exchanged with methionine-free media (RPMI, Thermo) containing AHA for 30 min. After treatment, media was removed, and cells were fixed and permeabilized. The AHA reaction cocktail was added to cells to detect AHA signaling as a marker of protein synthesis, and cells were co-labeled with Hoechst stain. Cells were visualized for fluorescent signal and total fluorescent intensity quantified in a ring around the nuclei using the Cell Insight CX7 HCX Platform.

### Respirometry and Hydrogen Peroxide Assay

Oroboros high-resolution respirometry was performed on fresh whole-tissue homogenate using an Oroboros Oxygraph-2K (Innsbruck, Austria). Homogenization was performed by adding 50 mg of frozen LV to Mir05 buffer (MgCl_2_-6H_2_O 3 mM, K^+^MES 105 mM, taurine 20 mM, KH_2_PO_4_ 10 mM, Hepes 20 mM, d-sucrose 110 mM, and fatty acid-free BSA 1 g/l). After baseline respiration was established, 5ͰmM pyruvate, 2ͰmM malate, and 10ͰmM glutamate (PMG) were added to assess respiratory capacity through complex I, followed by ADP (2 mM) titrations to stimulate respiration. Next, 10 μM cytochrome C was added to evaluate mitochondrial integrity, followed by 10 mM succinate to assess the combined activity of Complex I + II, then 0.5 μM carbonyl cyanide 3-chlorophenylhydrazone (CCCP) to uncouple the mitochondrial membrane and induce maximal respiration. Hydrogen peroxide levels were assessed from these samples using Amplex Red Assay (Thermo, #A22188) according to manufacturer instructions.

### Statistical Analysis

Data were expressed as mean ± standard error in all figures. P ≤ 0.05 was considered significant for all comparisons. One-way ANOVA with Dunnett’s multiple comparisons test was used to all compare groups except Figure 8, in which student’s t-test was used. All statistical analysis was completed using Graphpad Prism 8 software (San Diego, CA).

## Results

### Metformin pretreatment protects cardiac EF and is dependent on AMPK signaling

To test the cardioprotective effects of metformin in CA, mice were pretreated for 2-weeks with metformin before the CA procedure (Figure 1) and compared to untreated arrest mice. There were no significant differences between groups to baseline animal characteristics, including the ratio of female mice (Table 1), body weight (Table 1), or baseline EF (Figure 2A) in these mice. Twenty-four hours after arrest, untreated arrest mice had significantly lower EF than untreated sham mice (Sham: 59.5±1.7%; arrest: 41.1±2.7%, p<0.0001, Figure 2A), as expected based on our previous description of this model (26). Metformin pretreatment significantly protected EF measured 24 hours after CA (Arrest+Met: 51.6±2.6%, p<0.01 vs. Arrest, Figure 2A). There were no changes to postoperative body temperature, time to resuscitation, or time to extubation (Table 1). Sham mice pretreated with metformin had no change to baseline characteristics or EF 24 hours after the procedure. These baseline and control measures found no confounding variables.

**Table 1.** Surgical data for arrest mice. There are no significant changes to body weight, body temperature at time of extubation, time to return of spontaneous circulation (ROSC), time to extubation, or time to kidney reperfusion between groups when compared to the arrest group. Data are presented as mean ± SEM unless otherwise noted. Statistical comparisons by one-way ANOVA with Dunnett’s multiple comparison test.

|  | Sham<br>(n=20) | Arrest<br>(n=26) | Sham<br>Metformin<br>(n=8) | Arrest<br>Metformin<br>(n=21) | Arrest<br>AICAR<br>(n=9) | Arrest<br>Metformin+<br>compound C<br>(n=8) |
| --- | --- | --- | --- | --- | --- | --- |
| Age (d) | 57.55±0.68 | 57.60±0.59 | 58.75±0.40 | 58.80±0.48 | 58.44±0.50 | 58.75±0.37 |
| # Female / Total<br>Mice (percentage) | 11/20<br>(55%) | 12/26<br>(46%) | 5/8<br>(63%) | 12/21<br>(57%) | 4/9<br>(44%) | 4/8<br>(50%) |
| 8-wk Body Wt (g) | 23.19±0.83 | 22.97±0.62 | 21.05±0.88 | 21.77±0.78 | 22.59±1.1 | 22.47±1.23 |
| Temp at<br>Extubation (°C) | 35.86±0.15 | 35.53±0.20 | 35.96±0.18 | 35.95±0.12 | 35.79±0.19 | 36.21±0.11 |
| ROSC (min) | - | 1.31±0.09 | - | 1.43±0.15 | 1.44±0.24 | 1.23±0.12 |
| Time to<br>Extubation (min) | - | 22.31±0.66 | - | 23.64±0.32 | 22.06±0.90 | 24.06±0.52 |
| Time to Kidney<br>Reperfusion (min) | - | 21.3±0.90 | - | 21.79±0.57 | 20.43±1.01 | 24.70±0.64 |

**Figure 2.**
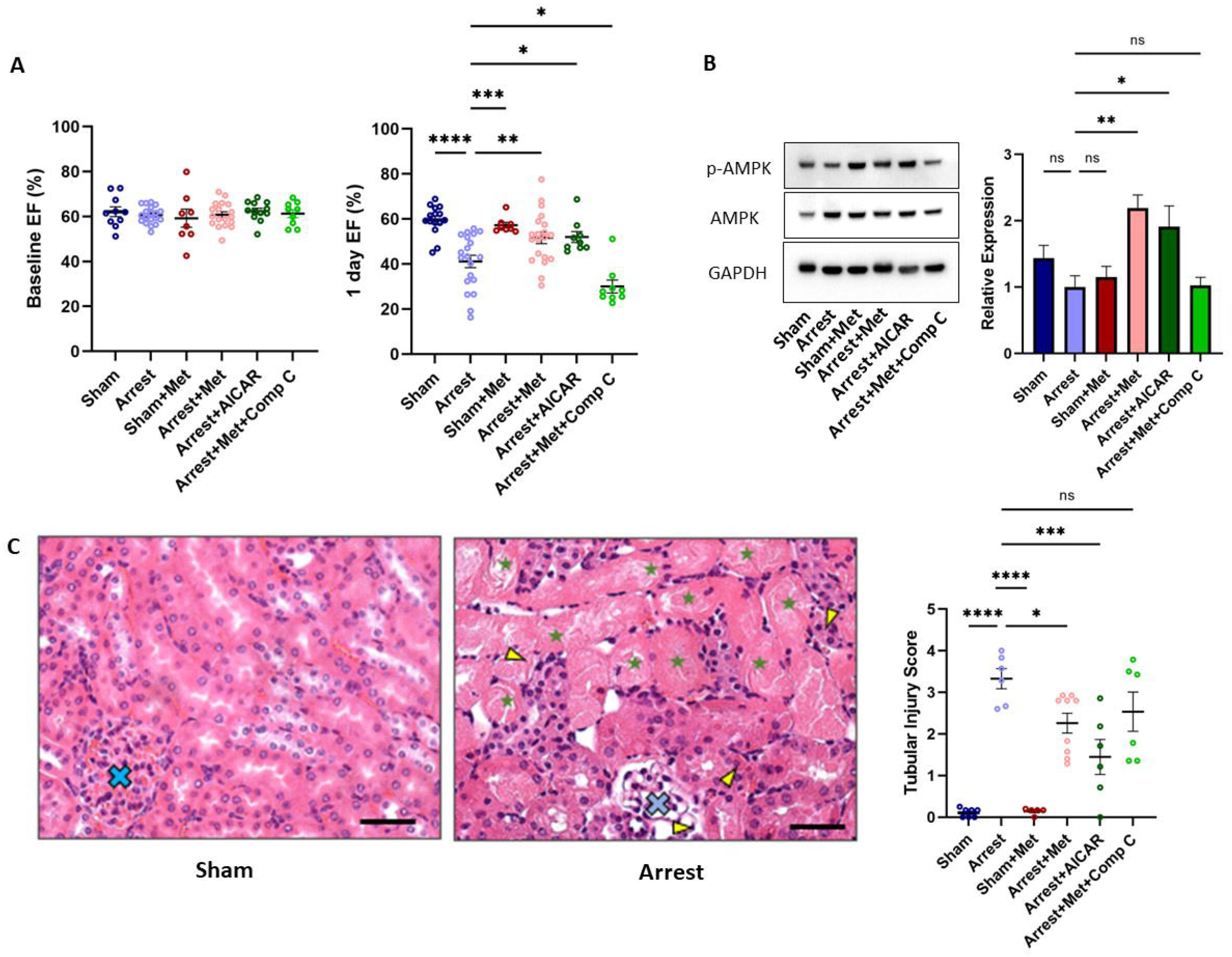
Metformin and AICAR treated mice have improved ejection fraction (EF), elevated cardiac p-AMPK/AMPK, and less kidney damage than untreated mice one day after cardiac arrest (CA). A) Baseline (left) and one day EF (right) among treatment groups. Circles represent individual mice. B) Representative Western blot analysis of p-AMPK (Thr172), AMPK, and GAPDH for each treatment group and quantification of Western comparing p-AMPK to total AMPK (n=6/group). Full blots are provided in Supplemental Figure 1. C) Representative histologic sections from untreated sham and untreated arrest mice demonstrating proteinaceous casts in renal tubules (green stars) and infiltrates (yellow arrow heads) with glomeruli marked (blue X’s). Scale bar = 50 μm) as well as quantification of tubular injury score between groups. Data are expressed as mean ± SEM. P-values: *< 0.05, **< 0.01, ***< 0.001, ****< 0.0001 by one-way ANOVA with Dunnett’s multiple comparison test. EF, ejection fraction.

Metformin has multiple potential modes of action (30), with AMPK activation being dominant in the literature. Because of AMPK’s role in cardiac resilience (25), we tested whether direct AMPK activation was necessary and sufficient for the observed enhanced EF. Cohorts of mice were treated for 2-weeks with the AMPK activator AICAR (31) or metformin and compound C, an established AMPK inhibitor (12). Both groups underwent two weeks of intraperitoneal (IP) injections before CA and were evaluated by echocardiography at baseline and twenty-four hours after arrest. No changes were observed in baseline EF (Figure 2A) or baseline characteristics, including age, % female, or body weight compared to the arrest group (Table 1). Like the metformin results, AICAR pretreated arrest mice had significantly improved EF compared to untreated arrest mice at 24 hours post-CA (Arrest+AICAR: 52.0±2.4%, p<0.05 vs. arrest, Figure 2A). Compound C not only prevented the beneficial effects of metformin on cardiac EF but also caused significantly reduced EF (Arrest+Met+Comp C: 30.0±2.9%) when compared to the untreated arrest group (p<0.05). The AICAR pretreatment phenocopies metformin pretreatment, and compound C blocks the benefit of metformin, suggesting that AMPK activation is necessary and sufficient for the metformin-mediated protection of cardiac function after CA.

To determine AMPK activation status, we assessed phosphorylation of threonine-172 of the AMPKα subunit in the myocardium (32). 24 hours after the CA procedure, LVs from sham and arrest groups (n=6/group) were assessed for p-AMPK, total AMPK, and glyceraldehyde 3-phosphate dehydrogenase (GAPDH) protein expression and were normalized to the arrest group (Figure 2B, Supplemental Figure 1). p-AMPK/AMPK was not significantly changed between Sham (1.44±0.47), Arrest (1.00±0.17), and Sham+Met (1.15±0.16) groups. p-AMPK/AMPK was elevated in the Arrest+Met mice (2.19±0.19, p<0.01 vs. Arrest) and Arrest+AICAR mice (1.91±0.31, p<0.05 vs. arrest) but unchanged in the Arrest+Met+Comp C mice (1.024±0.12). Total AMPK was unchanged between the treatment groups (Supplemental Figure 1). The strongest activation of AMPK was with metformin or AICAR plus arrest, suggesting a synergistic effect of drug and arrest on activation at this time point.

### Metformin pretreatment protects against kidney injury and is dependent on AMPK signaling

The effects of whole-body ischemia/reperfusion injury can be detected in peripheral tissues, particularly in the kidney (26). Untreated arrest mice have significant kidney damage by tubular injury scoring compared to untreated sham mice at one-day post-CA (sham: 0.11±0.04; arrest: 3.33±0.24, p<0.0001, Figure 2C). Metformin pretreated sham mice have no evidence of damage by tubular injury scores (0.14±0.04, p<0.0001 vs. arrest). Metformin pretreated arrest mice had significantly lower tubular injury scores (2.26±0.24, p<0.05) than the arrest group. AICAR pretreated arrest mice had lower injury scores (1.45±0.42, p<0.001) than arrest mice, while Metformin + compound C mice had no protection evident by score (2.54±0.47). To test peripheral markers as a proxy for tubular injury scores, serum markers of kidney damage, creatinine and BUN, were also assessed in each group (Supplemental Figure 2). Serum creatinine was significantly elevated in the arrest mice compared to sham (sham: 0.36±0.04 mg/dL; arrest: 1.49±0.14 mg/dL, p<0.0001). There was no significant change to metformin pretreated arrest mice (0.97±0.21 mg/dL), though significant changes were found in the AICAR arrest (0.67±0.25, p<0.01) and Metformin + compound C mice (0.85±0.07, p<0.05) compared to sham. BUN was elevated in arrest mice compared to sham (sham: 26.6±5.2 mg/dL; arrest: 153.9±28.4 mg/dL, p<0.001) but was unchanged between arrest mice and Metformin pretreated arrest (121.2±28.0 mg/dL), AICAR arrest (85.7±8.9), and Metformin + compound C mice (135.8±18.6). No changes were found when comparing the time to kidney reperfusion between the untreated arrest and arrest treatment groups by Doppler flow analysis of the renal artery (Table 1). Taken together, systemic activation of AMPK reduces histological markers of CA-induced kidney damage, but 24-hour serum markers show limited sensitivity to this protection.

### Metformin causes broad transcriptome changes in sham and arrest mice

To better understand the signaling changes caused by metformin therapy, transcriptomic analysis was performed on LVs from Sham, Arrest, Sham+Met, and Arrest+Met groups (n=6/group) by microarray. Differentially expressed genes were identified for three comparisons: Arrest vs. Sham, Sham+Met vs. Sham, and Arrest+Met vs. Arrest (p<0.05, no fold restriction, Supplemental Figure 3). Pathway analysis was then performed on these transcriptome changes through Ingenuity Pathway Analysis. The top 15 most significantly changed pathways are included in Figure 3, along with a z-score to predict the upregulation or downregulation of each pathway. Prominent pathways include changes to autophagy, AMPK signaling, unfolded protein response, nuclear factor-erythroid 2-related factor 2 (NRF2) mediated oxidative stress, and eukaryotic initiation factor 2 (EIF2) signaling. We performed initial confirmation studies for these pathways and focused on changes detectable at 24 hours. When no supporting data were found, we advanced other pathway studies. When support was identified, additional studies were done for confirmation.

**Figure 3.**
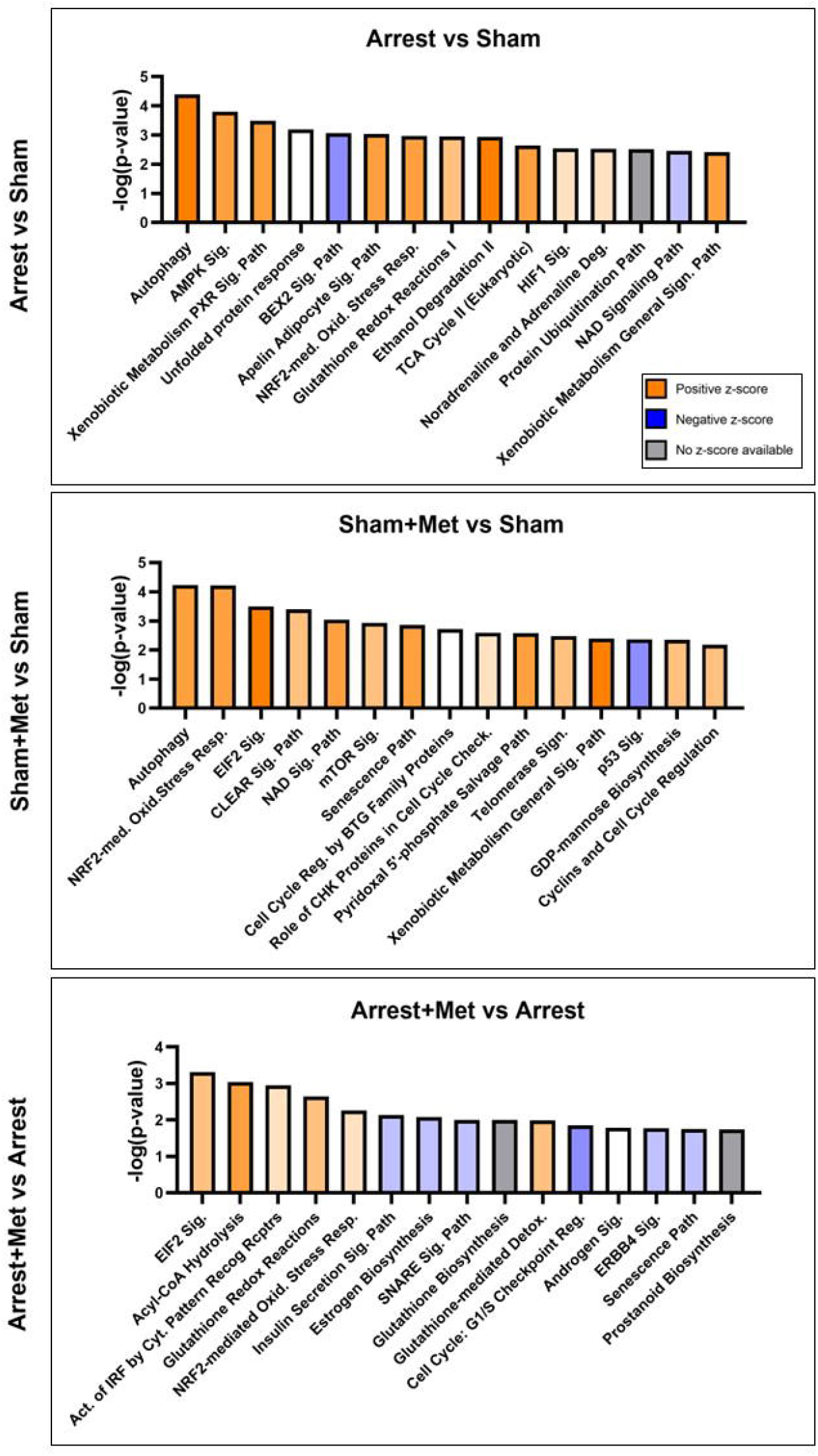
Pathway Analysis Comparing Treatment Groups. Ingenuity Pathway Analysis was used to predict signaling pathways between Arrest and Sham mice (top), Sham+Metformin (Sham+Met) and Sham mice (middle), and Arrest+Metformin (Arrest+Met) to Arrest mice (bottom) based on microarray data collected from left ventricles one day after arrest (n=6/group). Bar-graphs include significance values for pathway changes (-log(p-value)) and are colored according to z-score to predict directionality of changes. Differentially expressed genes were included in the analysis when p<0.05 based on microarray data. Principal component analysis and volcano plots are included in Supplemental Figure 2.

### Metformin affects mitochondrial morphology and markers of autophagy

Interestingly, the autophagy pathway shows the most significant changes between Arrest and Sham and Sham+Met and Sham, but not between Arrest+Met and Arrest. Because metformin has been reported to increase cardiac mitophagy in cardiomyopathy (33,34) and autophagy was the top hit in two of the sample comparisons, we evaluated protein expression for markers associated with autophagosome formation, including p62/Sequestosome 1, a cargo receptor associated with degradation of ubiquitinated proteins (35), and microtubule-associated protein light chain 3 (LC3) processing (Figure 4). p62 expression (normalized to GAPDH) was not significantly changed between groups. The relative levels of microtubule-associated protein light chain 3 (LC3), specifically levels of uncleaved (LC3-I) and cleaved (LC3-II) forms, were also monitored as an indicator of changes in autophagy initiation (36). Interestingly, the LC3-II to LC3-I ratio was significantly increased in untreated arrest mice (1.91±0.15) compared to sham (1.00±0.18, p<0.01). In contrast, the metformin-pretreated arrest mice had significantly reduced LC3-II/LC3-I (1.17±0.17) when compared to untreated arrest mice (p<0.05). LC3-I and LC3-II expression levels (normalized to GAPDH) were not significantly changed between groups (data not shown). The normalization of LC3-II/LC3-I may be interpreted as metformin restoration of mitophagy; however, ischemia/reperfusion is well known to induce mitochondrial damage and mitophagy (37), and reduction in mitochondrial damage could yield a similar change in LC3 ratios.

**Figure 4.**
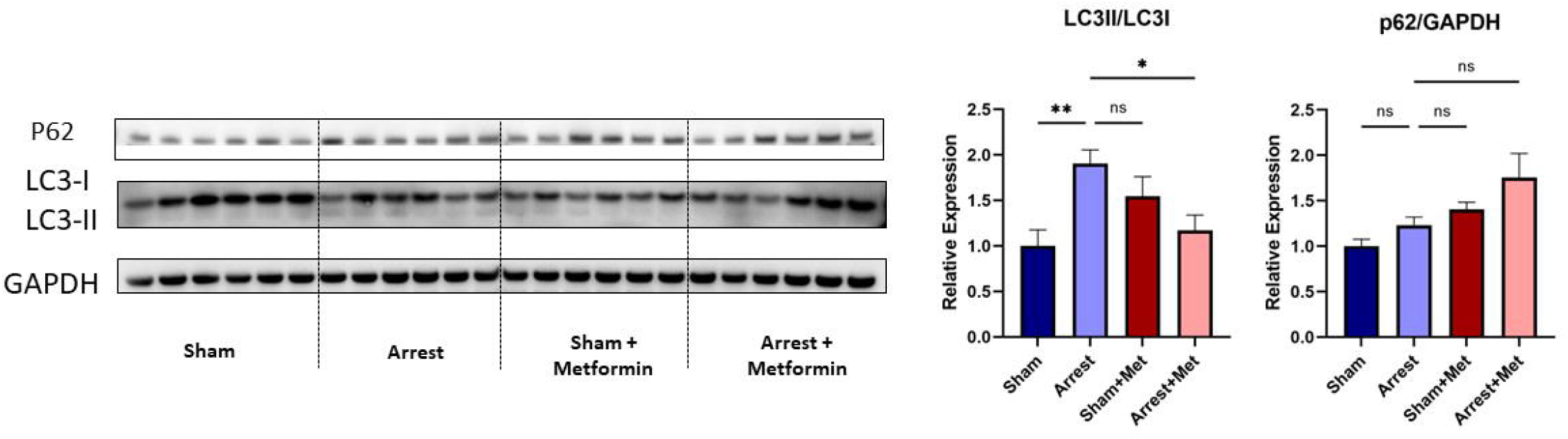
Western Blot Analysis of Autophagy Markers. Western blot images of autophagy-related proteins downstream of AMPK including p62, LC3 (I/II), and housekeeping protein (GAPDH) expression in Sham, Arrest, Sham+Metformin (Sham+Met), and Arrest+Metformin (Arrest+Met) mice (n=6/group). Data are expressed as mean ± SEM. P-values: *< 0.05, **< 0.01 by one-way ANOVA with with Dunnett’s multiple comparison.

To assess mitochondrial damage, we used electron microscopy to examine mitochondrial structural changes among hearts from untreated sham, untreated arrest, and metformin-pretreated arrest mice (Figure 5A). The untreated arrest mice showed a decrease in mitochondrial perimeter and area (perimeter: 3.06±0.06 μm; area: 0.60±0.02 μm^2^) when compared to sham (perimeter: 3.81±0.09 μm; area: 0.87±0.04 μm^2^, p<0.0001 for both measures) in cardiomyocytes (Figure 5B). Myocardium from metformin-pretreated arrest mice showed a modest but significant increase in mitochondrial perimeter and area (perimeter: 3.33±0.07 μm; area: 0.70±0.03 μm^2^) when compared to that of untreated arrest mice (p<0.05 for both measures). Both the untreated arrest mice (0.76±0.001 μm, p<0.0001) and metformin-pretreated arrest mouse heart (0.75±0.01 μm; p<0.0001) had more circular mitochondria than the untreated sham mouse (0.69±0.01 μm). These data demonstrate modest changes in mitochondrial morphology at 24 hours post-CA, with normalization of mitochondrial area and perimeter but not circularity in metformin-pretreated arrest mice compared to untreated arrest mice, which may be consistent with diminished mitochondrial damage or normalization of membrane dynamics.

**Figure 5.**
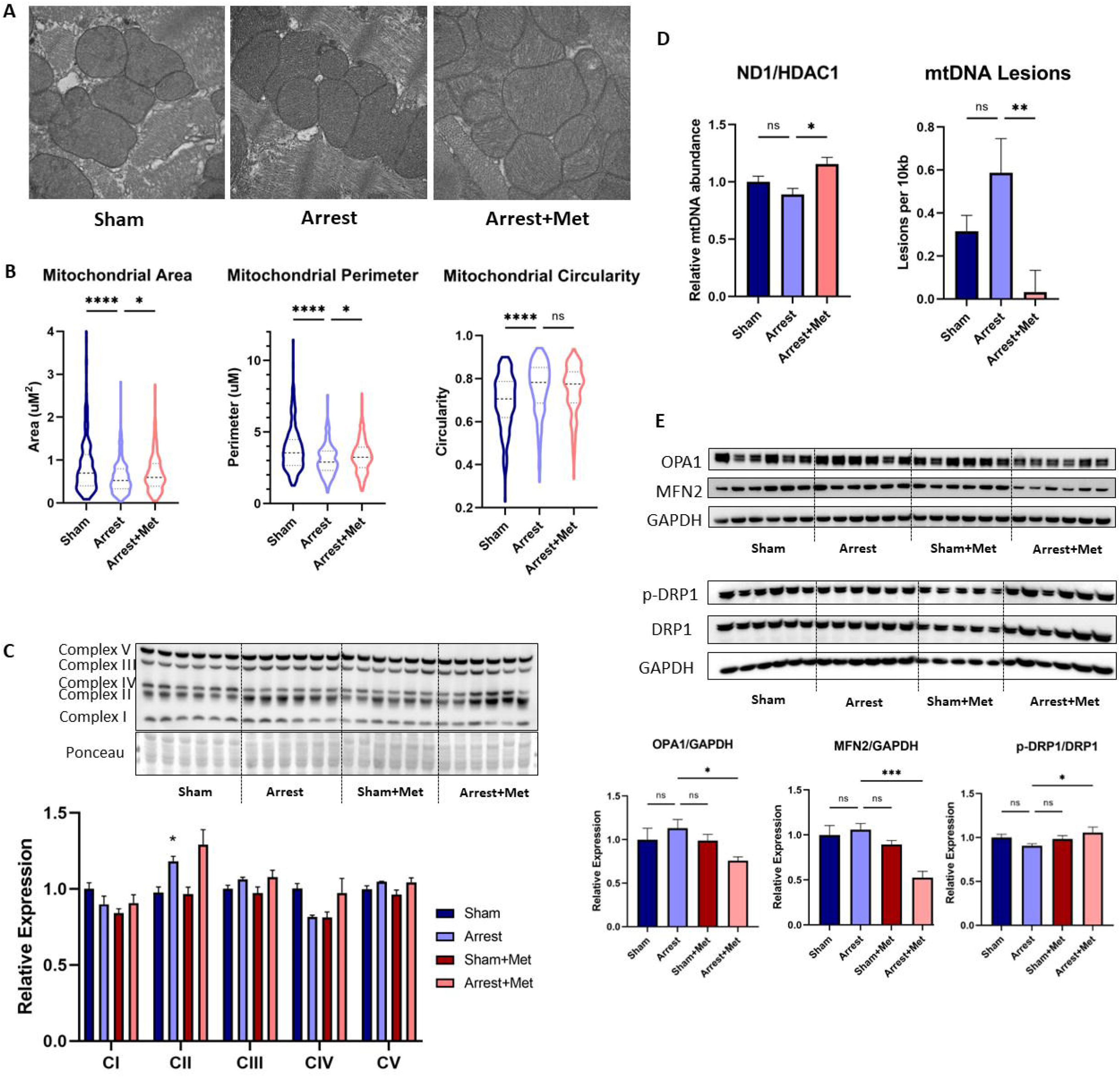
Metformin pretreatment affects mitochondrial size, mtDNA damage, and mitochondrial fission/fusion proteins after cardiac arrest (CA). A) Representative electron microscope images of intrafibrillar mitochondria in Sham, Arrest, and Arrest+Metformin (Arrest+Met) mice one day after surgery (40,000x magnification) B) Violin plot showing mitochondrial perimeter, area, and circularity (n=50 per group). C) Western blot of representative oxidative phosphorylation complex subunits (CI-CV) and bar graph of densitometry (n=6/group, normalized to ponceau stain). Quantitation shown in bar graphs with indicated normalization. D) Bar graph of mtDNA copy number assessment by qPCR quantitation of total DNA preparationsof sham (n=5), arrest (n=4), and arrest metformin (n=7) mice. E) Western blot images of markers of mitochondrial fission and fusion (MFN2, OPA1, p-DRP1 (ser616), DRP1) and housekeeping protein (GAPDH) in Sham (n=6), Arrest (n=6), Sham+Met (n=5-6), and Arrest+Met (n=6) mice. B) C). E) Data are expressed as mean ± SEM. P-values: *< 0.05, **< 0.01, ***< 0.001, ***<0.0001 by one-way ANOVA with with Dunnett’s multiple comparison. DRP, dynamin-related protein1; MFN2, mitofusin 2; OPA1, dynamin-like 120 kDa protein, mitochondria1.

Because of the change in mitochondrial architecture, we also looked at markers of mitochondrial biogenesis and damage in the LVs to explain the protection by metformin. First, representative proteins of mitochondrial respiratory complexes were assessed from LV extracts taken 24 hours after arrest from untreated sham, untreated arrest, metformin-pretreated sham, and metformin-pretreated arrest mice (Figure 5C). There was a mild but statically significant elevation in a subunit of complex II in the metformin-pretreated arrest mice (1.29±0.10) compared to untreated sham mice (0.98±0.04, p<0.01); but otherwise, relative expression of representative proteins was unchanged across the groups. Consistent with these results, we found no significant differences in Oroboros measurement of mitochondrial oxygen consumption in heart homogenates in sham and arrest cohorts (Supplemental Figure 4A). After sequential stimulation with PMG, ADP, Cytochrome C, succinate, and FCCP, we found no significant differences in oxygen consumption rates (OCR). Prior work shows that mitochondrial dysfunction induced by ischemia-reperfusion injury resolved in less than 24 hours (38,39), so this result only informs that the respiratory defect is not protracted in metformin+arrest heart.

### Metformin pretreatment protects against mtDNA damage after arrest

We next examined mtDNA relative abundance and damage in the cardiac tissue from these groups (Figure 5D). Unlike in failed hearts (40), the arrest group did not show a statistically significant decrease in mtDNA levels at 24 hours post-CA, consistent with the evidence above that mitochondria are not being dramatically removed by autophagy. The metformin-pretreated arrest mice had slightly higher relative mitochondrial DNA (mtDNA) levels (1.16±0.06) than untreated arrest mice (0.89±0.05, p<0.05) but were not significantly different from the sham group (1.00±0.05). Notably, the metformin-pretreated arrest mice had significantly less mtDNA damage in a long extension assay (0.03±0.1 lesions/10kb mtDNA) than untreated arrest mice (0.59±0.16 lesions, p<0.05). There was no statistical difference between sham and arrest+met groups. The small but significant increases in mtDNA and complex II in the metformin-pretreated arrest mice could be consistent with an increase in mitochondrial biogenesis, but the combined significant improvement in mitochondrial morphology and mtDNA integrity suggests an improvement in overall mitochondrial quality, decreased damage, and are in line with the changes in autophagy markers.

### Metformin pretreatment affects markers of mitochondrial dynamics

To understand the alterations in mitochondrial morphology better, we examined the relative levels of several proteins involved in establishing mitochondrial shape (Figure 5E). Mitofusin 2 (MFN2), a mitochondrial outer membrane GTPase involved in fusion, had reduced expression in metformin-pretreated arrest mice (Arrest+Met: 0.53±0.07) compared to all other groups (Sham: 1.00±0.10, p<0.01; Arrest: 1.06±0.07, p<0.001; Sham+Met: 0.89±0.04, p<0.05). OPA1, also a dynamin-related GTPase that resides in the inner membrane to perform fusogenic functions (41), followed a similar overall trend with significantly reduced levels in metformin-pretreated arrest mice (0.75±0.04) when compared to untreated arrest mice (1.13±0.10, p<0.05). In contrast, dynamin-related protein 1 (DRP1), whose activity is regulated by its phosphorylation at Ser-616 (42), showed no significant change in p-DRP1/DRP1 between sham and arrest groups. Metformin pretreatment appeared to improve p-DRP1/DRP1 post-arrest modestly, but the effect was not statistically significant, likely because the decline in p-DRP in untreated arrest hearts was not significant. Indeed, while mitochondrial perimeter and area are increasing with metformin-pretreatment in 24-hour post-arrest hearts, markers of fusion and fission are not significantly changed by arrest (relative to sham), suggesting that these markers are not the driver of those morphological changes. In fact, the MFN2 and OPA1 decrease and p-DRP1/DRP1 increase by metformin pretreatment in arrested mice (relative to arrest mice) are moving in the opposite direction from what would be expected to correct mitochondrial morphology. We suggest that other mitochondrial quality control processes contribute to mitochondrial morphological rescue.

### Metformin pretreatment protects against changes to markers of protein synthesis

Interestingly, the MFN2 decrease is consistent with the activation of autophagy (43,44), where it is ubiquitinated and targeted for proteolysis. Metformin has been reported to impact heart autophagy through AMPK signaling (33,34). To test engagement of autophagy in the CA model, LVs collected 24 hours after arrest from untreated sham, untreated arrest, metformin-pretreated sham, and metformin-pretreated arrest mice were evaluated for markers of autophagy by Western blot analysis (Figure 6). First, we evaluated mTOR activation as a crucial negative regulator of autophagy (45). As a positive downstream marker of mTOR activity, S6 ribosomal protein (S6) phosphorylation at Ser-240/244 (p-S6) was assessed (Figure 6A). Both the p-S6/GAPDH and p-S6/S6 showed that the mTOR pathway activity was highest in the arrest + metformin. We suggest that mTOR is potentially highly activated in arrest+metformin hearts at 24 hours post-arrest because mTOR activity inhibits autophagy, the concept that AMPK activation and subsequent mTOR inhibition to increase autophagy is insufficient to explain the survival benefit of metformin.

**Figure 6.**
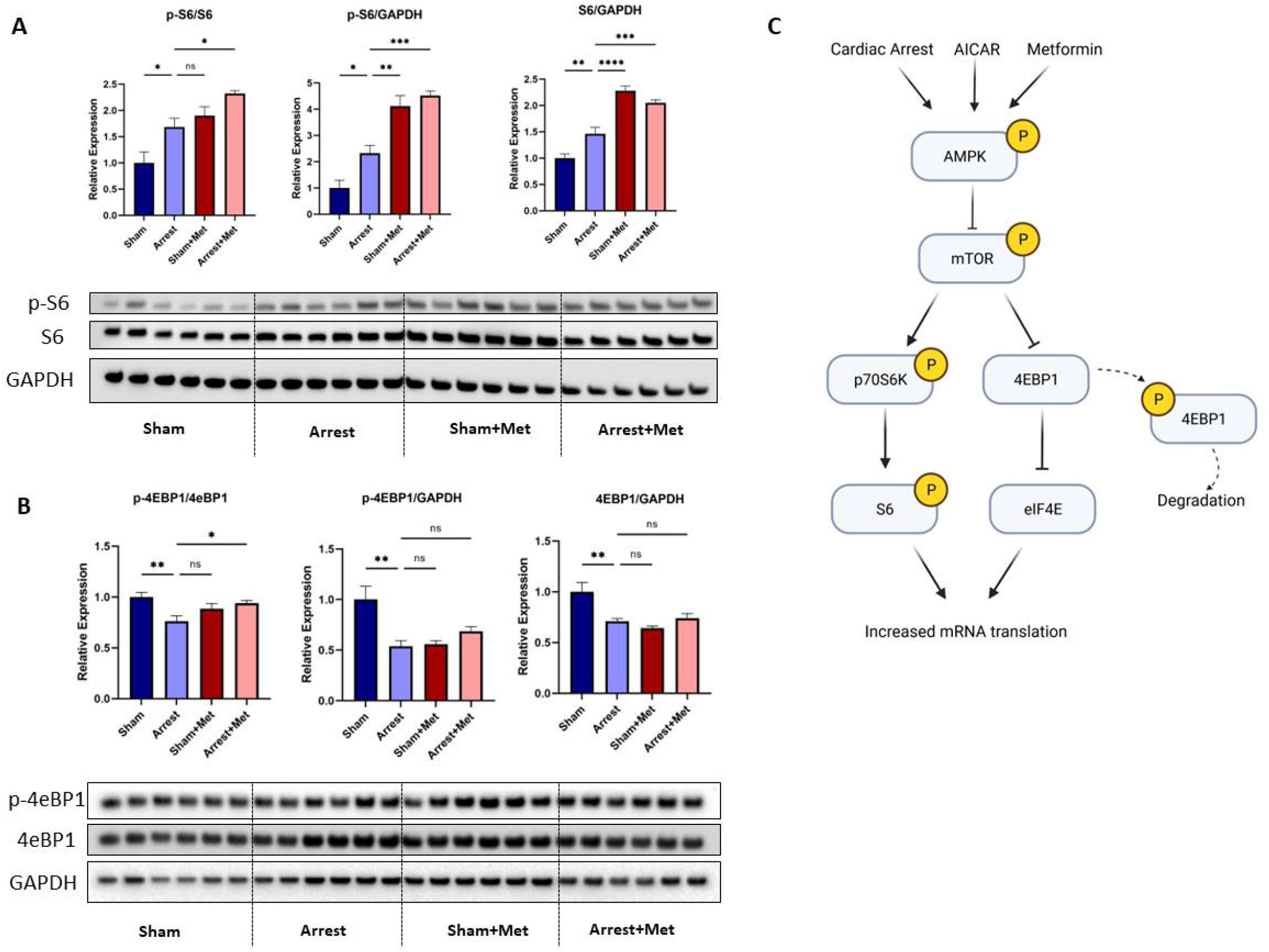
Western Blot Analysis of Protein Synthesis Markers. A) Western blot images of S6 and its phosphorylated form at Ser240/244 and housekeeping protein (GAPDH) expression in Sham, Arrest, Sham+Metformin (Sham+Met), and Arrest+Metformin (Arrest+Met) mice (n=6/group). B) Western blot images of 4eBP1 and its phosphorylated form at Thr37/46 as well as GAPDH. C) Cartoon representation of p-S6 and p-4EBP1 as regulators of mRNA translation downstream of AMPK and mTOR signaling. Data are expressed as mean ± SEM. P-values: *< 0.05, **< 0.01, ***< 0.001 by one-way ANOVA with with Dunnett’s multiple comparison test. 4eBP1, eukaryotic translation initiation factor 4e-binding protein 1; mTOR, mechanistic target of rapamycin; S6, ribosomal protein S6.

EIF2 signaling was the most significantly altered pathway between the Arrest+Met and Arrest groups (Figure 3) and is activated by mTOR (46,47). EIF2 signaling is commonly referred to as a marker of the endoplasmic stress downstream of protein kinase RNA-like ER kinase (PERK) activation (48). In that role, EIF2 activation promotes the expression of activating transcription factors (ATFs), including ATF4, an integrated stress response regulator known to be activated by metformin (49,50). However, we did not find significant changes in ATF4 expression between our treatment groups (Supplemental Figure 5). The mRNA changes driving EIF2 signaling are overwhelmingly related to ribosomal proteins (Supplemental Figure 5B), leading us to explore markers of protein synthesis.

As both the EIF2 pathway and p-S6 activation increase mRNA translation, we next performed an additional Western blot analysis of eukaryotic translation initiation factor 4E-binding protein 1 (4EBP1). 4EBP1 is a repressor of mRNA translation (51), and when phosphorylated at Thr37/36, 4EBP1 releases eukaryotic translation initiation factor 4E (eIF4E) and is degraded (Figure 6C). eIF4E. proceeds to increase protein synthesis. We found that the ratio of p-4EBP1/4EBP1 was decreased in arrest mice (0.76±0.05) when compared to sham (1.00±0.05, p<0.01; Figure 6B). The Sham+Met (0.89±0.05) group was unchanged from Arrest; however, the Arrest+Met group (0.94±0.03) was significantly higher than Arrest (p<0.05). Sham mice had higher p-4EB1/GAPDH (1.00±0.13) than Arrest mice (0.54±0.05, p<0.01), while there was no change between the Arrest mice and the Sham+Met (0.56±0.04) or the Arrest+Met (0.69±0.05) groups. Similarly, Sham mice had higher 4EBP1/GAPDH (1.00±0.09) vs. Arrest (0.71±0.03, p<0.01), but there was no change between Arrest mice and Sham+Met (0.64±0.02) or Arrest+Met (0.74±0). The elevation of p-S6, p-4EBP1, and activation of the EIF2 pathway strongly implicate mTOR activation in promoting translation that persists at 24 hours post-arrest.

### AICAR pretreatment protects against H/R changes to protein synthesis in a cell model

To quantify protein synthesis changes, we replicated the hypoxic insult of CA in cell culture by using AC16 cells (Figure 7A). Cells were pretreated overnight with AICAR to mimic AMPK activation specifically. Cells underwent normoxia or hypoxia (1% oxygen in Esumi buffer) for 4 hours before reoxygenation with and without AICAR pretreatment. Using these conditions, we found that p-S6/S6 was decreased in the untreated H/R cells (0.78±0.01) compared to untreated normoxia (1.00±0.06, p<0.01) and unchanged compared to AICAR normoxia (0.91±0.02) but was significantly higher in the AICAR H/R cells (0.93±0.05, p<0.05). Similar changes were found in the p-S6/GAPDH and S6/GAPDH comparisons. Finally, p-4EBP1/4EBP1 was decreased in untreated H/R cells (0.72±0.01) compared to untreated normoxia (1.00±0.08, p<0.05) and unchanged compared to AICAR normoxia (0.83±0.03) and AICAR H/R cells (0.87±0.06). However, p-4EBP1 was significantly lower in the untreated H/R cells (0.56±0.01) compared to all other groups (untreated normoxia: 1.00±0.04, p<0.001, AICAR normoxia; 0.74±0.04, p<0.01, AICAR H/R: 0.74±0.04, p<0.01). There was no change in the percent of dead cells after reoxygenation in any cell groups (Figure 7C).

**Figure 7.**
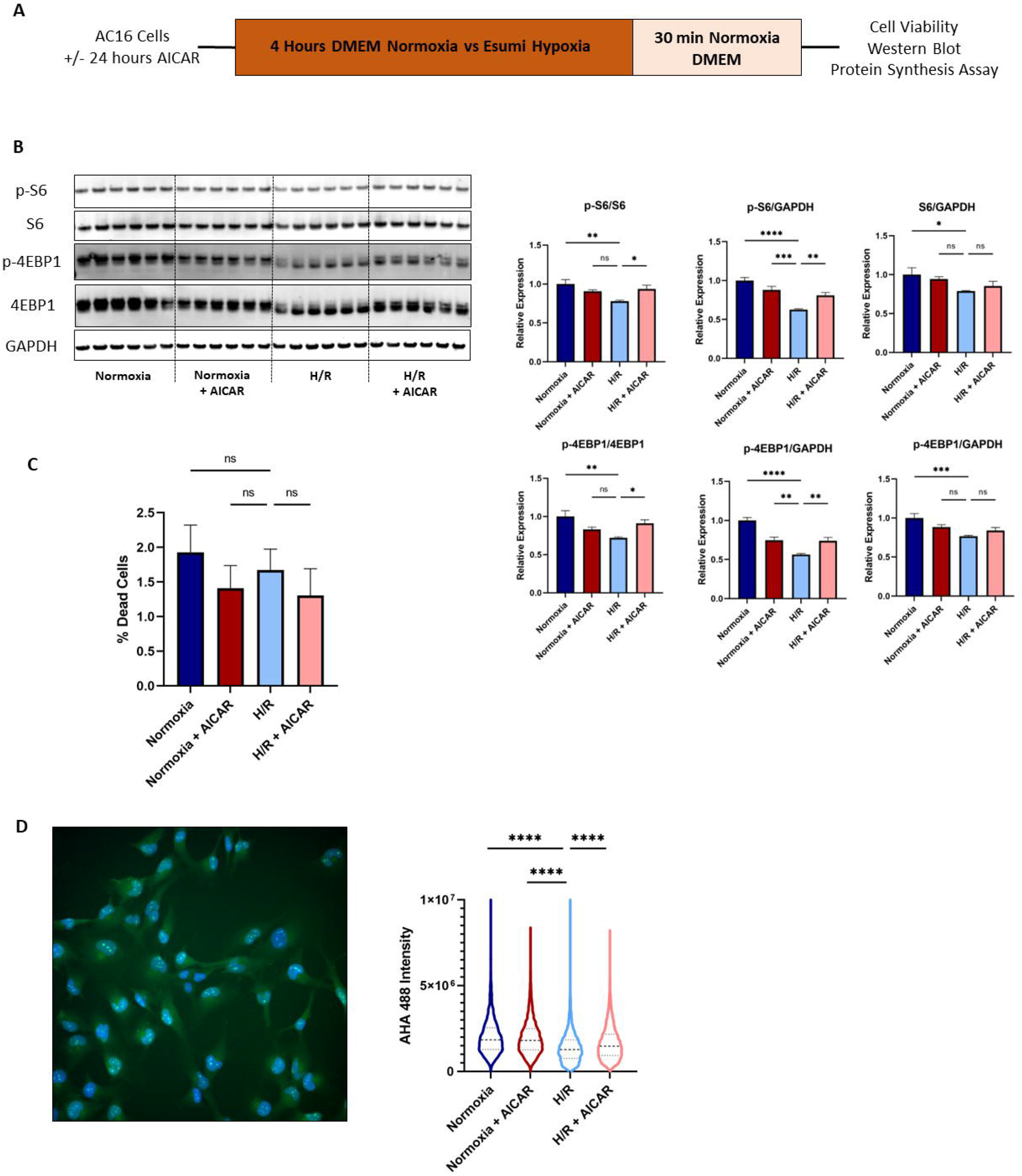
AICAR pretreatment protects against protein synthesis changes in AC16 cells after hypoxia/reoxygenation (H/R). A) Time course of AC16 cells with and without 24 hours of AICAR treatment (1.25 mM). B) Western Blot images of of p-S6 (Ser240/244), total S6, p-4EBP1 (Thr37/46), total 4EBP1, and GAPDH expression from cells 30 min after H/R (n=6/group). C) Quantification of cell death 30-min after H/R. D) Representative image of AHA 488 Protein Synthesis assay demonstrating AHA 488 signal (green) and Hoechst nuclear stain (blue) in AC16 cells 30 min after H/R and quantification of AHA 488 signal intensity surrounding the nuclei after H/R. P-values: *< 0.05, **< 0.01, ***< 0.001 by one-way ANOVA with with Dunnett’s multiple comparison test.

Protein synthesis was quantified for each cell group by measuring the fluorescent intensity of labeled AHA given immediately after reoxygenation (n=3000-5000 cells/group). The fluorescent intensity was lower in the untreated H/R cells (1397713±12864) compared to untreated normoxia (2021451±13931, p<0.0001) and AICAR normoxia cells (1969457±16463, p<0.0001; Figure 7D). The AICAR H/R cells had significantly higher intensity (1651628±16842) compared to the untreated H/R cells (p<0.0001). These *in vitro* data suggest that activation of AMPK would restore protein translation disrupted by ischemia/reperfusion injury in heart.

### Metformin does not improve outcomes as a rescue therapy

To test metformin’s efficacy as rescue therapy in our model, metformin was given directly into the LV in a cohort of mice (n=7) at the time of resuscitation (1,250 μg/kg). Compared to the arrest mice described in Figure 2, there was no change to EF one day after CA (arrest rescue metformin: 42.9±2.4%, Figure 8A). Similarly, there was no change in tubular injury score (2.54±0.423, Figure 8A), creatinine (1.2±0.25, Supplemental Figure 6), or BUN (155.3±14.2) when compared to untreated arrest mice. Baseline EF (61.1±2.1) was not significantly different from the arrest mice. These data are consistent with the notion that metabolic adaptation is required for metformin protection after CA.

**Figure 8.**
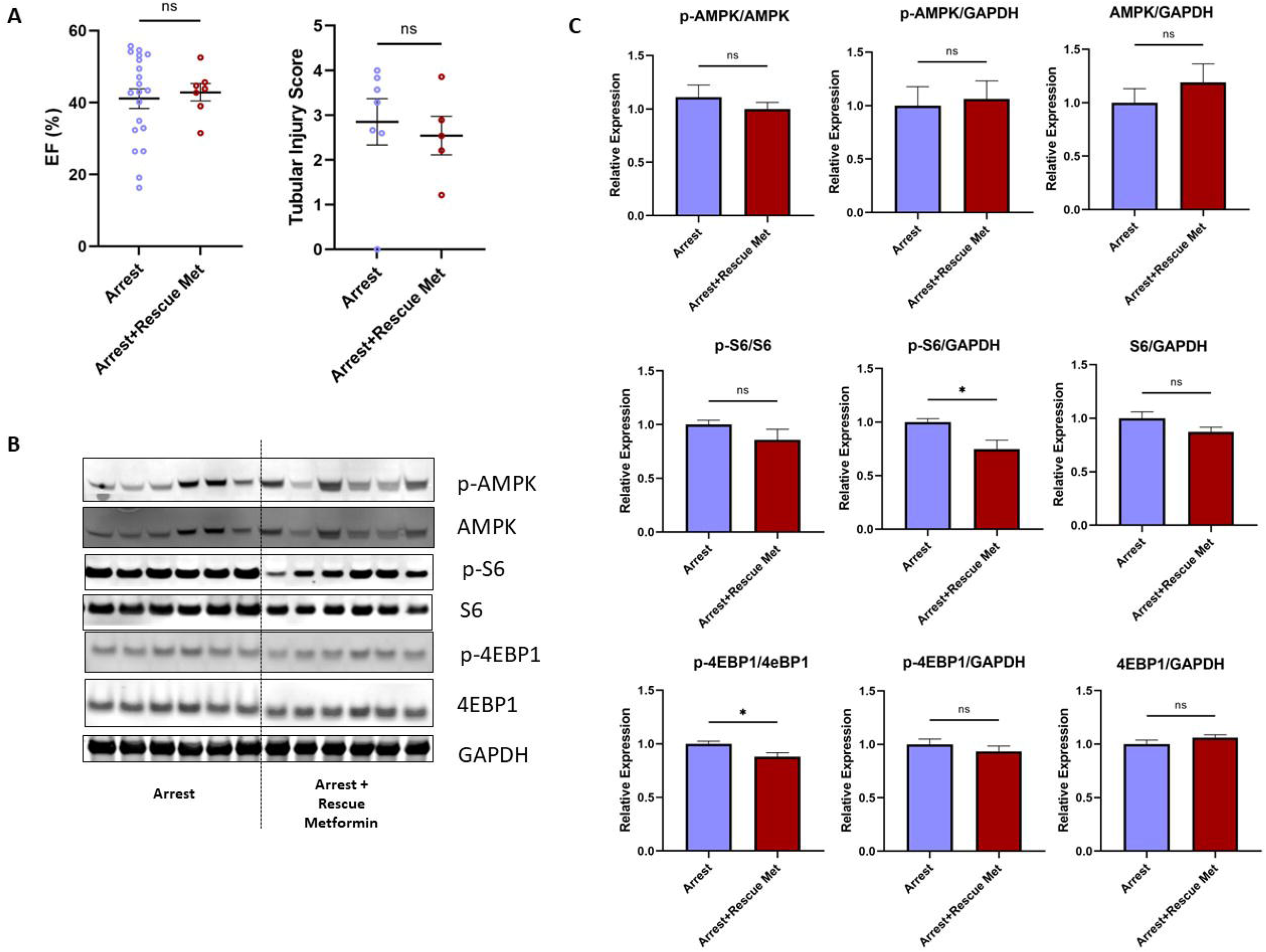
Metformin does not protect cardiac or renal function as a rescue therapy following arrest. A) When given concomitantly with epinephrine, intravenous metformin (rescue met, n=7) did not demonstrate any change to ejection fraction (EF) when compared to untreated arrest mice (n=20; arrest data also presented in Figure 2) or to kidney damage as measured by tubular injury score B) Western Blot images of of p-AMPK (Thr172), total AMPK, p-S6 (Ser240/244), total S6, p-4EBP1 (Thr37/46), total 4EBP1, and GAPDH expression between arrest and arrest + rescue metformin groups (n=6/group) C) Quantification of Western blot changes between groups, *p<0.05 by student’s t-test.

Hearts from these mice were collected, and Western blots performed comparing untreated arrest vs. arrest with rescue metformin. There was no change in p-AMPK/AMPK, p-AMPK/GAPDH, or AMPK/GAPDH between treatment groups (Figure 8B&C). p-S6/S6 and S6/GAPDH were unchanged between groups, though there was a reduction in p-S6/GAPDH in the arrest + rescue metformin cohort (0.75±0.08) when compared to untreated arrest mice (1.00±0.03, p<0.05). There was a mild reduction in p-4EBP1/4EBP1 in the arrest + rescue metformin mice (0.88±0.04) when compared to untreated arrest mice (1.00±0.02, p<0.05), but p-4EBP1/GAPDH and 4EBP1/GAPDH were unchanged. These differences in magnitude and duration of activation of these pathways in this model may account for the failure as a rescue therapy.

## Discussion

Our group recently performed a retrospective analysis of diabetic CA patients with and without metformin therapy before arrest and found that metformin improved markers of cardiac and renal damage acutely after CA (9). This exciting result prompted the search for reports of preclinical models to identify the molecular pathways involved. Of particular relevance are studies of a 9-min rat asphyxial model. Metformin pretreatment has neurologic and 7-day survival benefits but focuses on neuronal markers without measures of cardiac and renal outcomes (21). More recently, a 10-min asphyxial rat model with post-CA treatment demonstrated improved 3-day survival and neurological function, involving normalization of brain activity without activation of AMPK, but also did not examine cardiac and renal function (22). Our current data suggest that metformin-mediated cardiac protection may also promote survival. Indeed, the lack of AMPK activation in the 2-hours post-treatment brain could be consistent with protection arising from other metformin activities. Unfortunately, acute metformin rescue was unsuccessful in our myocardial stunning model, so potential confounders in interpretation among these studies remain.

Metformin pretreatment has previously decreased scar size in reversible coronary artery ligation (24,52–56) and *ex vivo* whole-heart ischemia-reperfusion (24,57). However, studies of metformin’s effects on cardiac function have primarily utilized ischemia-reperfusion injury models with ischemic periods of 25-30 minutes. The long ischemic duration in these models involves substantial cardiomyocyte necrosis. Much of metformin’s beneficial effect has been attributed to reducing mPTP-mediated cell death (58). In contrast, our data demonstrate metformin’s protection of *in vivo* EF in a CA model (Figure 1) that features an 8-minute ischemia period without evidence of cardiac cell death (26). We, therefore, set out to identify intracellular processes that adapt the cardiomyocyte to maintain function rather than alter cell survival.

Metformin pretreatment has pleiotropic effects involving many molecular pathways, including reducing oxidative stress, inhibiting apoptosis, inhibiting complex I, and activating AMPK signaling (20,24,59,60). In our CA mouse model, we find that AMPK activation is necessary and sufficient to enact metformin’s cardioprotective effects. Mice pretreated with AICAR, an AMP-mimetic known to activate AMPK (50), had preserved EF one day after CA compared to the untreated arrest mice (Figure 2A). Conversely, mice pretreated with metformin with the AMPK inhibitor compound C showed no improvement in EF one day after arrest compared to the untreated arrest mice. AICAR conferred similar renal protection as measured by tubular injury score, and the protective effects of metformin on the kidney were negated by compound C (Figure 2C). Interestingly, we did not find evidence of p-AMPK/AMPK changes between the arrest and sham groups by Western blot (Figure 2B), despite indicated AMPK changes in transcriptome pathway analysis (Figure 3). This phosphorylation status of AMPK may have recovered at the time point evaluated (24 hours after arrest), while the continued effects of metformin and AICAR are observed.

Pathway analysis was performed on microarray data collected from LVs of Sham, Arrest, Sham+Met, and Arrest+Met mice. A comparison of arrest to sham mice (Figure 3) implicates autophagy and AMPK signaling in our model. Given the evidence of metformin’s AMPK-dependent effects (Figure 2), it should be expected that metformin therapy causes similar pathway changes in the sham mouse or the untreated arrest model. Several signaling changes were observed when comparing untreated arrest to untreated sham mice (Figure 3, top) when comparing the metformin-treated sham to untreated sham mice (Figure 3, middle). These shared pathways include activation of autophagy, NRF-2 mediated oxidative stress response, NAD signaling, and xenobiotic metabolism signaling. Given this overlap, it is unsurprising that signaling pathways are less profound when comparing the metformin pretreated arrest and untreated arrest mice (Figure 3, bottom). Interestingly, the strongest hit in the Arrest+Met mice compared to the arrest animals was EIF2 signaling, which may suggest an increased ability to support the initiation of translation for protein synthesis after CA.

We focused on transcriptional pathways that were altered at the time of cardiac assessment, which showed overt cardiac dysfunction (Figure 1A). Autophagy, AMPK signaling, unfolded protein response, Nrf2-mediated oxidative stress, and EIF2 signaling show differences in transcriptional signatures between Sham, Arrest, and Arrest+Met at 24 hours post-arrest. The absence of changes in gene expression between arrest and metformin arrest may be due to the preservation of cardiac function being upstream of those alterations. In the process, we observed whether given pathway markers being tested showed activation or normalization, i.e., regression to sham levels. For autophagy, the pathway was activated in Arrest vs. Sham and Sham+Met vs. Sham but *not* in Arrest+Met vs.Arrest, suggesting that the pathway wasn’t accelerated at the time of analysis. Normalization of mitochondrial function, mitochondrial morphology, LC3-II/I ratios, and hyperphosphorylation of S6, which is inhibitory for autophagy (61), are consistent with the non-activation of autophagy. Although mtDNA lesions are decreased, which could be compatible with accelerated autophagy, the increase in total mtDNA levels and mitochondrial Complex II may indicate some compensatory mitochondrial biogenesis. The significant decreases (i.e., not normalization) of OPA1 and MFN2 and increase in p-DRP1 levels in Arrest+Met relative to Arrest alone (Figure 5E) would suggest a change in mitochondrial dynamics, which has previously been implicated as a major regulator in cardiac arrest outcomes (62). However, we find that mitochondrial area and perimeter are increasing mitochondrial size (Figure 5B) instead of the expected shift to a more fragmented shape due to changes in those proteins (41,63,64). We did not detect oxidative protein modification (Supplemental Figure 4), which is known to be primarily mitochondrial proteins damaged at early timepoints (38,39), suggesting that oxidant activating the Nrf2 signaling pathway may have been resolved by 24 hours post-CA. This is in line with the normalization of mitochondrial function by 24 hours post-reperfusion described by others and the lack of differences in respiration described herein (Supplemental Figure 4A).

The hyperactivation of p-S6 also showed an accelerated response at 24 hours post-CA in Arrest+Met compared to Arrest alone samples. Initial tests of ATF4 activation downstream of S6K activation did not support the unfolded protein response’s involvement at this time. We speculated that p-S6 was not only indicating that AMPK stimulated activity inhibited mTOR, which activated p70S6K, but also the downstream activation of translation (Figure 6C). We also show alterations to the phosphorylation status of the translation repressor protein 4EBP1 in the untreated arrest mice, which is normalized in the arrest mice pretreated with metformin (Figure 6B). Phosphorylated 4EBP1 is dissociated from the translation initiation factor eIF4E (Figure 6C), allowing eIF4E to initiate translation (65).

To confirm changes in protein synthesis, we performed H/R studies of AC16 cells. We found that 4 hours of ischemia in these cells was enough to insult to drive Western blot markers without causing cell death (Figure 7A, C). Cell homogenates in the untreated H/R groups have elevated p-AMPK/GAPDH compared to normoxia and AICAR-treated normoxia cells (Figure 7B). However, we did not find superactivation of p-AMPK in the AICAR pretreated H/R cells. p-S6/S6 was significantly decreased in the untreated H/R cells compared to normoxia but preserved in the AICAR pretreated H/R group. Similarly, p-4EBP1/4EBP1 was reduced in the H/R mice compared to untreated normoxia cells but preserved in the AICAR pretreated group. The protective effects of AICAR on p-S6/S6 and p-4EBP1/4EBP1 were comparable to those seen by metformin in the animal model.

It should be noted that the p-S6/S6 is lower in the untreated H/R cells compared to the untreated normoxia cells, which is discordant with changes seen in the metformin-treated mice; however, the mouse model was complicated by epinephrine injection and surgical handling as well as delayed tissue collection (24 hours in mice, compared to 30 min in cells), which may explain the differences in p-S6/S6 results between the *in vivo* and *in vitro* systems. Importantly, we found that untreated H/R cells reduced protein translation compared to untreated normoxia cells, partially protected by AICAR pretreatment (Figure 7D). While these data suggest that AMPK activation can preserve translation in ischemia/reperfusion stressed cardiomyocytes, additional studies will be needed to establish to what extent such preservation contributes to cardiomyocyte resilience *in vivo*.

Finally, we performed a direct injection of metformin into the LV at the time of resuscitation to test the effects of metformin as rescue therapy. To minimize animal injury, we compared the rescue metformin cohort to the large arrest cohort described in Figure 2. However, we completed the analysis of EF and kidney changes in a blinded fashion. This small cohort of mice had no evidence of improved EF or kidney injury compared to the arrest group (Figure 8, Supplemental Figure 7). There has been some controversy concerning the use of metformin after cardiac insult. While several preclinical studies have found beneficial effects of metformin therapy when given at the reperfusion (24,56,59,66), this protection was not evident in a pig model of myocardial infarction (67). Our data suggest that while metformin as a pretreatment protects both the heart and the kidneys against ischemia-reperfusion injury, this protection depends on adaptive mechanisms before the insult and the nature of the insult. When used as rescue therapy, metformin did increase p-S6/S6 as seen in the metformin pretreatment group, but p-4EBP1/4EBP1 was significantly lower in the rescue metformin mice. This supports the notion that metformin’s support for mRNA translation after CA requires preconditioning rather than rescue therapy.

There are limitations to the results observed in this study. First, while metformin provided clear evidence of renal protection as measured by tubular injury (Figure 2C), these results are less clear when comparing serum markers of kidney injury, including creatinine and BUN. Further, we found unexpected protection against renal damage in the Arrest+Met+Comp C cohort compared to Arrest, which may suggest an additional signaling component in the kidney or varying degrees of AMPK activation between the heart and the kidney. However, these serum measurements depend on other physiologic factors, such as animal fluid status or peripheral tissue breakdown, which may complicate their interpretation. For this reason, we consider tubular injury score a superior marker of damage (68). This renal damage does not appear to depend on kidney reperfusion time (Table 1). However, this reperfusion time was measured by doppler tracing of the renal artery, which may not have the sensitivity to fully detect ischemic duration without contrast imaging (69). Regarding the rescue therapy, intravenous metformin was calculated based on pharmacologic rodent studies comparing oral to intravenous metformin bioavailability (70). The concentration was 10-times larger than an intravenous dose of metformin used as rescue therapy in a mouse infarction model (56), so it is unlikely that this concentration did not exert a significant effect.

In summary, we demonstrate that pretreatment with metformin protects against myocardial stunning and renal injury in a mouse model of CA. Our CA mouse involves cardiac injury without cardiac cell death, making it a more focused system to study metformin’s mechanism of action. Direct AMPK activation and inhibition studies confirmed that AMPK activation is necessary and sufficient for the cardiac and renal benefits observed with metformin treatment. While multiple molecular mechanisms downstream of AMPK activation may affect these outcomes, microarray data and Western blot changes suggest that preservation of mRNA translation may be a major mechanism underlying metformin’s effects. Preserved mRNA translation and protein synthesis were also found in a cell model of H/R when pretreated with AICAR. Unfortunately, these findings did not translate into use as rescue therapy, suggesting that metformin’s effects require a degree of adaptation to confer tissue protection.

## Acknowledgments

CR and BK were responsible for the conceptualization of these studies. CR designed the methodology, and CL, KR, and EG performed the investigation. Formal analyses were performed by CR, CL, TC, EG, and SSL. TC and DS completed visualization. CR, SSL, and BK wrote the manuscript. SSL and BK supervised the project, and BK provided resources for its completion. All authors reviewed the final manuscript and are responsible for its integrity. Some content was created with BioRender.

