## Supplemental Figures for "Metformin preconditioning protects against myocardial stunning and preserves mitochondrial dynamics and protein translation in a mouse model of cardiac arrest"

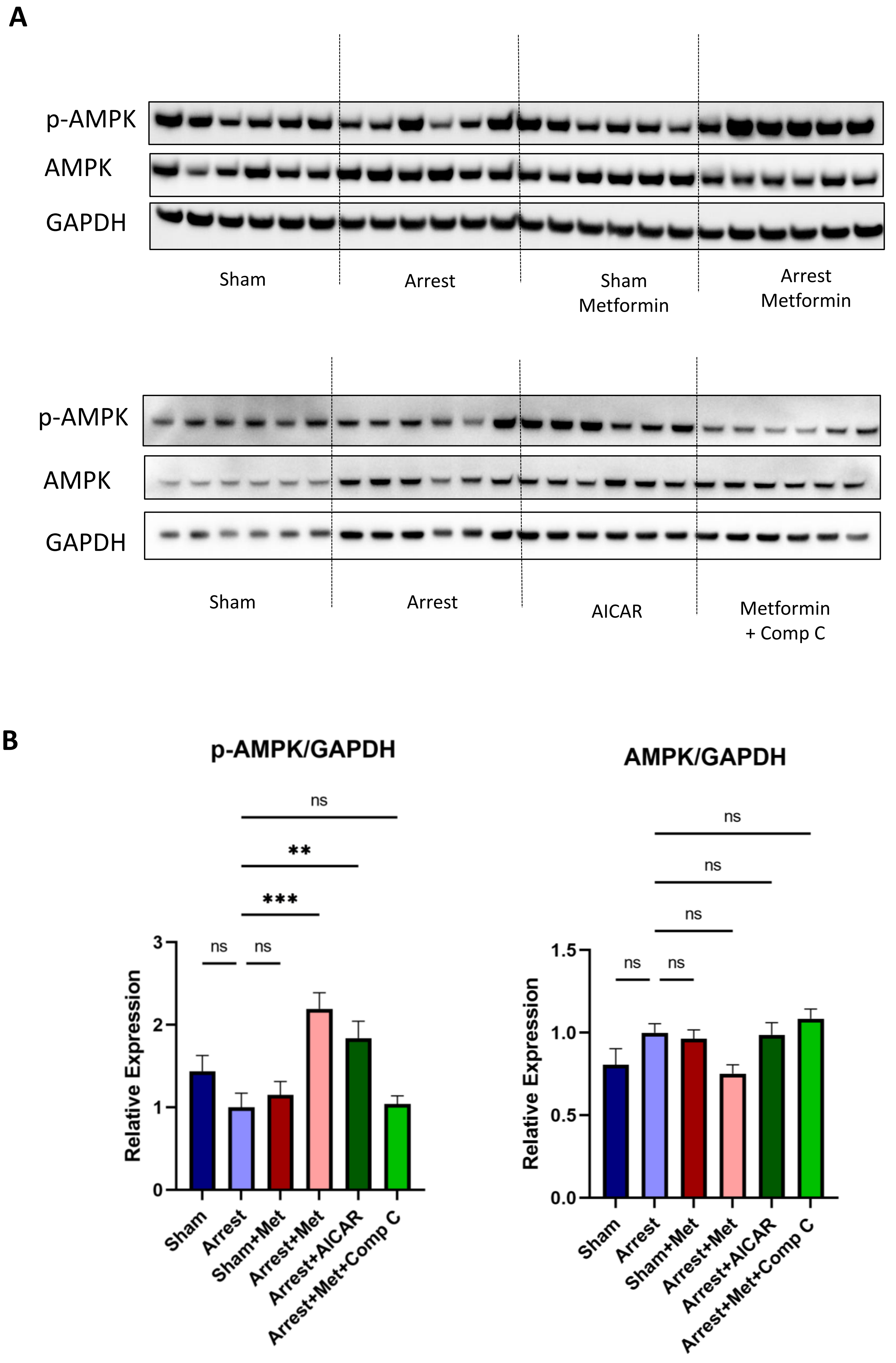


**Supplemental Figure 1. Western Blot Analysis of p-AMPK and AMPK.** A) Original blots of p-AMPK (Thr172), total AMPK, p-S6 (Ser240/244), and GAPDH for sham, arrest, sham+metformin, and arrest+metformin (top) as well as sham, arrest, arrest+AICAR, and arrest+metformin+compound C (bottom). B) Quantification of Western blots by densitometry. All samples were normalized to the arrest group and compared across blots.

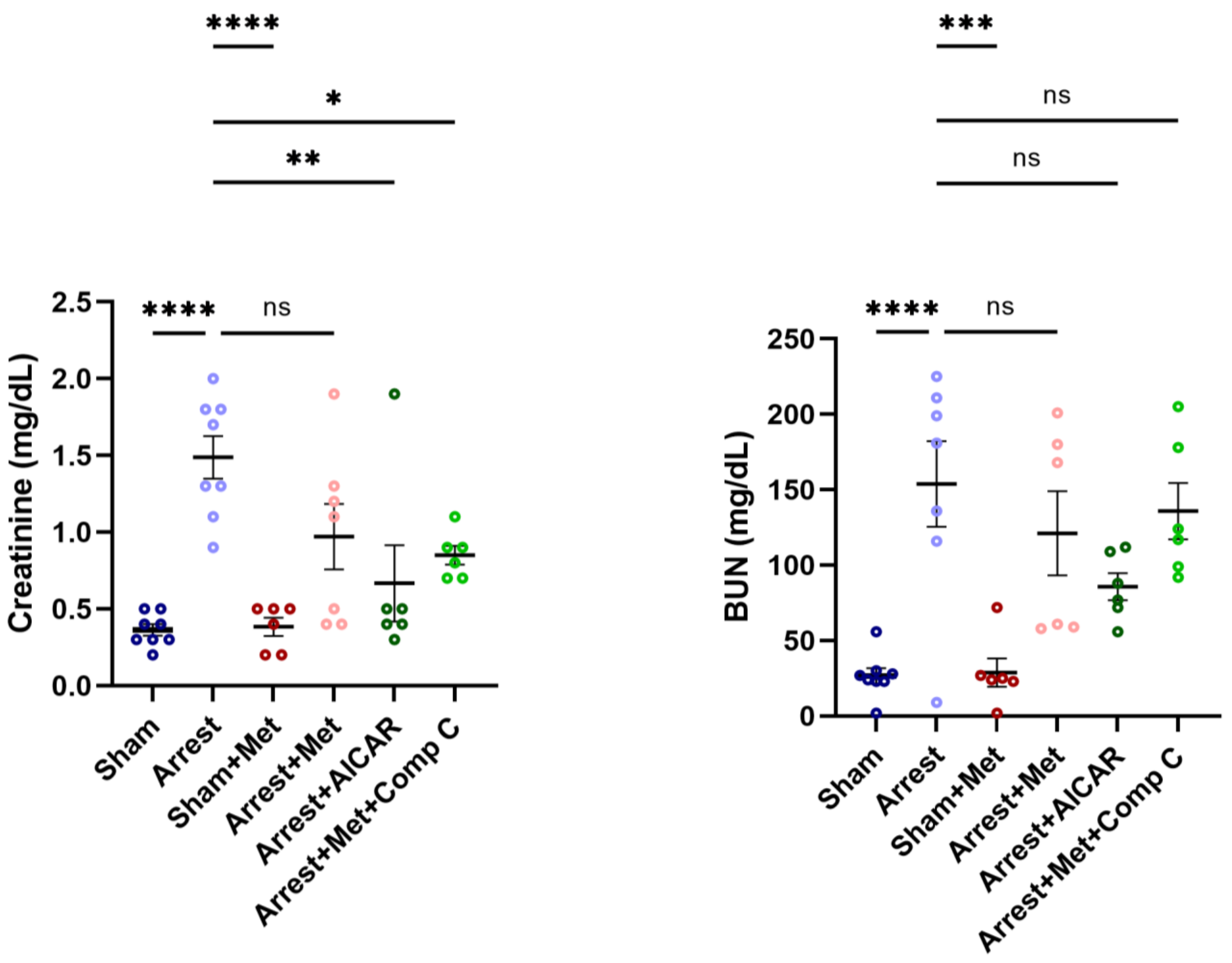


**Supplemental Figure 2. BUN and Creatinine in pretreatment groups.** Serum markers of kidney damage, including creatinine and blood urea nitrogen (BUN). Data are expressed as mean ± SEM. P-values: *< 0.05, **< 0.01, ***< 0.001 by one-way ANOVA with Dunnett's multiple comparison test.

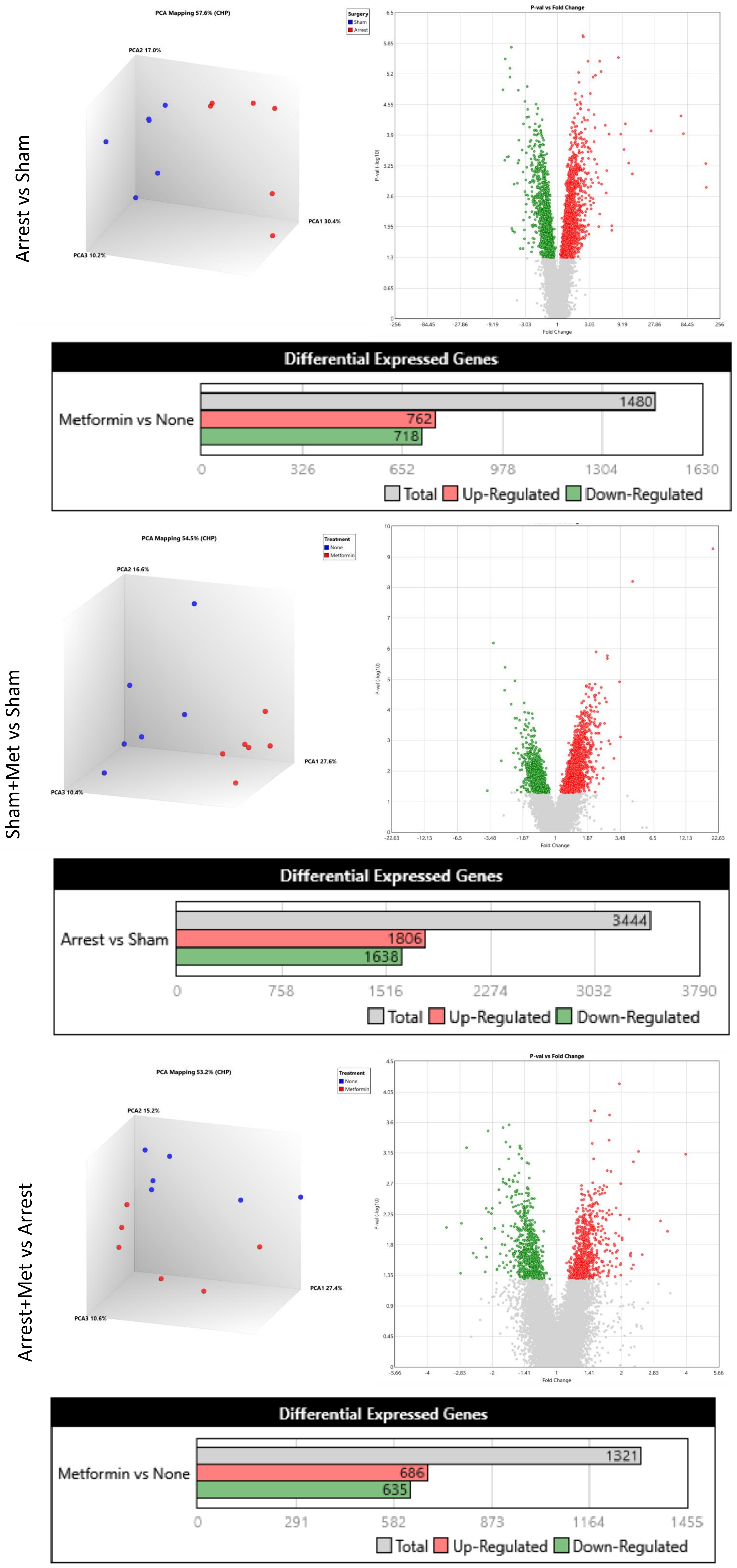


**Supplemental Figure 3. Microarray Data and Differentially Expressed Genes.** Microarray analysis was performed on Sham, Arrest, Sham+Metformin, and Arrest+Metformin mice (n=6/group). Principal component analyses (PCA) and volcano plots ($p<0.05$, no restriction on fold changes) were generated by Transcriptome Analysis Console. A summary of differentially expressed genes is included for each comparison group.

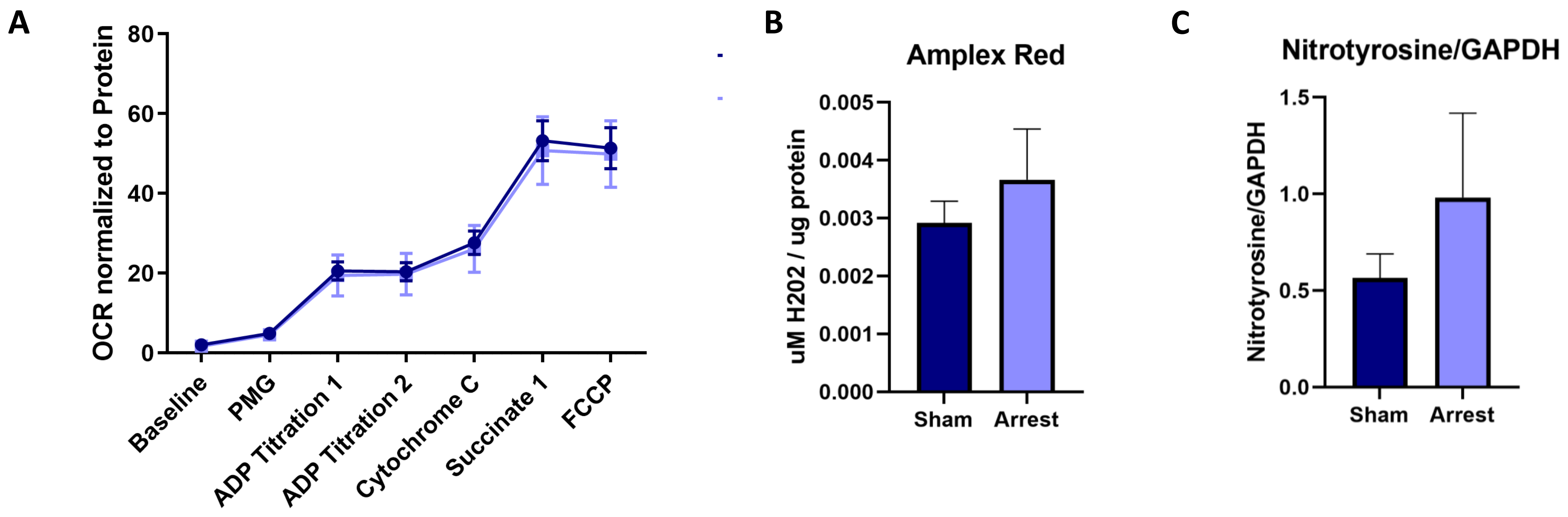


**Supplemental Figure 4. Oroboros oxygen consumption rate (OCR) and markers of oxidative stress.** A) There are no changes to OCR between Sham and Arrest groups 24 hours after arrest at baseline, complex I respiratory capacity after addition of pyruvate, malate, and glutamate (PMG,), ADP, complex I+II expression after cytochrome c, or uncoupled respiration after FCCP (n=4/group). B) There is no change to hydrogen peroxide accumulation as measured by amplex red assay (n=4-6/group). C) There is no change to protein nitrosylation by Western blot (n=4-6/group). Data are expressed as mean ± SEM and significance measured by student's t-test.

A

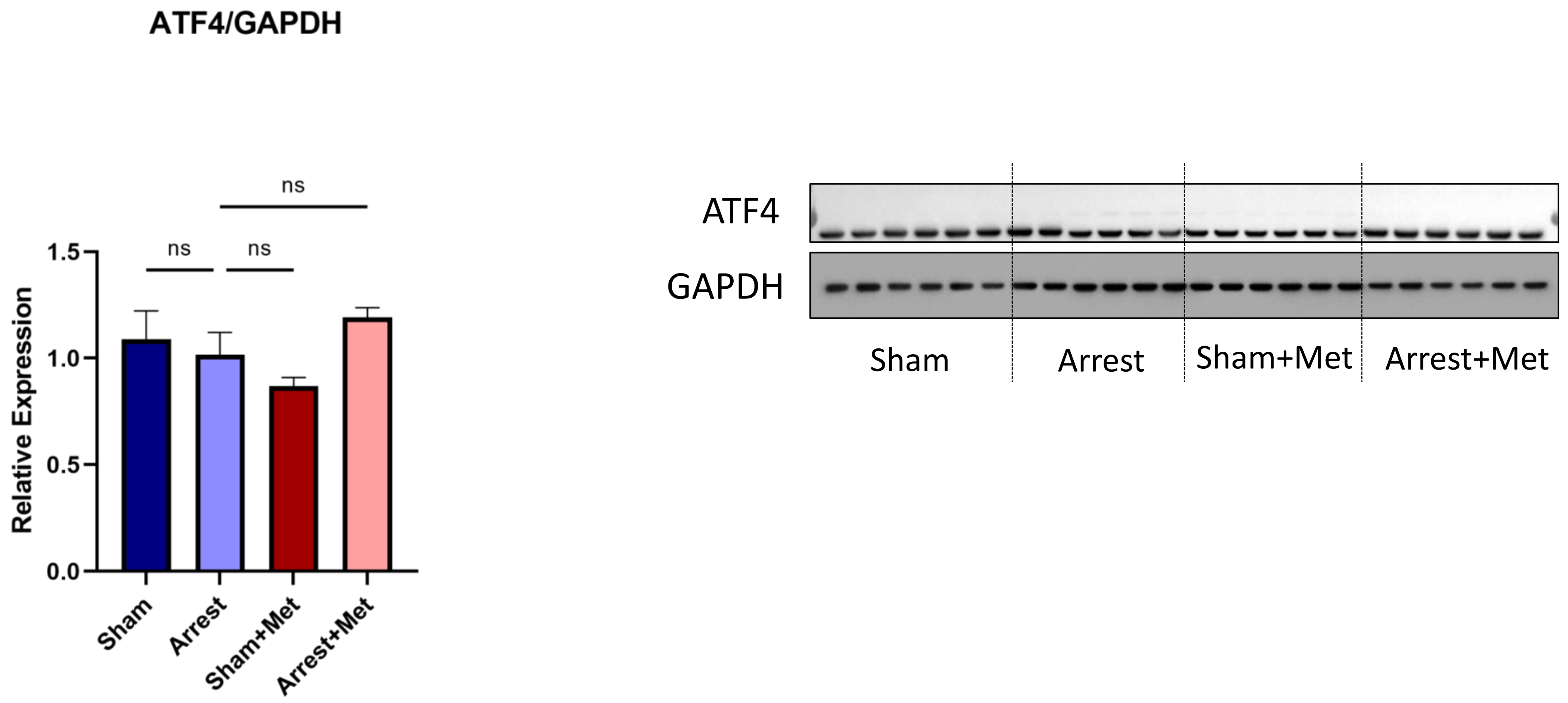


B

| Symbol | Entrez Gene Name | Affymetrix | Expr Fold Change (Arrest+Met vs Arrest) |
|---|---|---|---|
| ACTA1 | actin alpha 1, skeletal muscle | TC0800003161.mm.2 | -1.51 |
| PAIP1 | poly(A) binding protein interacting protein 1 | TC1300001346.mm.2 | -1.22 |
| PIK3C2G | phosphatidylinositol-4-phosphate 3-kinase catalytic subunit type 2 gamma | TC0600001647.mm.2 | -1.33 |
| PTBP1 | polypyrimidine tract binding protein 1 | TC1000000820.mm.2 | 1.64 |
| RPL3 | ribosomal protein L3 | TC1500001896.mm.2 | 1.58 |
| RPL6 | ribosomal protein L6 | TC0500001353.mm.2 | 1.23 |
| RPL10 | ribosomal protein L10 | TC0X00000704.mm.2 | 1.57 |
| RPL11 | ribosomal protein L11 | TC0400003779.mm.2 | 1.46 |
| RPL12 | ribosomal protein L12 | TC0200000600.mm.2 | 1.44 |
| RPL13 | ribosomal protein L13 | TC0800001508.mm.2 | 1.46 |
| RPL15 | ribosomal protein L15 | TC1400001500.mm.2 | 1.38 |
| RPL23 | ribosomal protein L23 | TC1100003655.mm.2 | 1.27 |
| RPL35 | ribosomal protein L35 | TC0200003433.mm.2 | 1.42 |
| RPL38 | ribosomal protein L38 | TC1100001874.mm.2 | 1.55 |
| RPL41 | ribosomal protein L41 | TC1000003151.mm.2 | 1.31 |
| RPL13A | ribosomal protein L13a | TC0700004648.mm.2 | 1.24 |
| RPS5 | ribosomal protein S5 | TC0700000151.mm.2 | 1.43 |
| RPS7 | ribosomal protein S7 | TC1200001568.mm.2 | 1.21 |
| RPS8 | ribosomal protein S8 | TC0400003362.mm.2 | 1.34 |
| RPS13 | ribosomal protein S13 | TC0700004155.mm.2 | 1.26 |
| RPS21 | ribosomal protein S21 | TC0200002785.mm.2 | 1.45 |
| RPS28 | ribosomal protein S28 | TC1700001879.mm.2 | 1.33 |
| RPSA | ribosomal protein SA | TC0900001595.mm.2 | 1.62 |
| RRAS2 | RAS related 2 | TC0700004132.mm.2 | 1.88 |

**Supplemental Figure 5. EIF2 Signaling Pathway in Arrest+Met vs Arrest Mice.** A) ATF4 expression is unchanged between treatment groups. Data are expressed as mean ± SEM. No significant changes by one-way ANOVA with with Dunnett’s multiple comparison test. B) Transcriptome changes between Arrest+Met and Arrest mice related to EIF2 signaling cascade as measured by microarray analysis and characterized by Ingenuity Pathway analysis.

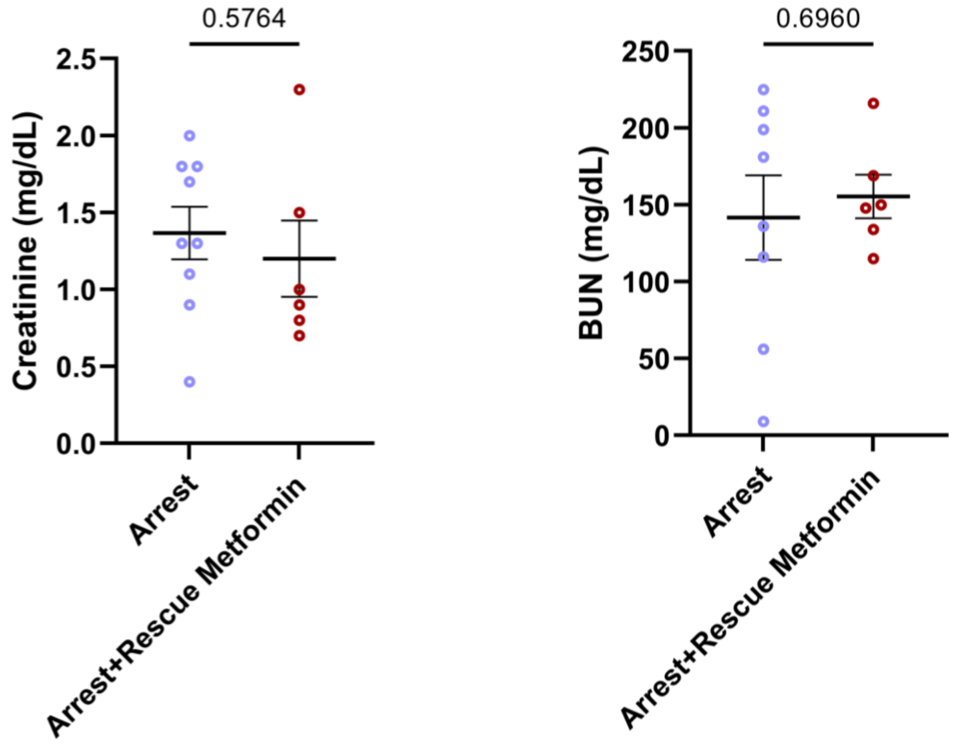


**Supplemental Figure 6. BUN and Creatinine data for rescue metformin.** Serum markers of kidney damage, including creatinine and blood urea nitrogen (BUN). Data are expressed as mean ± SEM and significance measured by student's t-test.
